# Identification of novel leaf rust seedling resistance loci in Iranian bread wheat germplasm using genome-wide association mapping

**DOI:** 10.1101/2022.02.24.481752

**Authors:** Saba Delfan, Mohammad Reza Bihamta, Seyed Taha Dadrezaei, Alireza Abbasi, Hadi Alipour, Jafargholi Imani, Karl-Heinz Kogel

## Abstract

Leaf or brown rust caused by *Puccinia triticina* Eriks. (*Pt*) is a major biotic constraint threatening bread wheat production worldwide. The continued evolution of new races of *Pt* necessitates a constant search for the identification of new resistance genes, or QTLs, to enhance the resistance durability of bread varieties. On a panel of 320 bread wheat accessions, we used a genome-wide association study (GWAS) technique to map loci associated with *Pt* resistance using single-nucleotide polymorphism markers (SNPs) generated by genotyping-by-sequencing (GBS). The panel was tested with five *Pt* races gathered from different regions of IRAN to identify loci associated with seedling resistance. After estimating genetic relatedness and population structure among accessions, GWAS discovered a total of 19 SNPs on chromosomes 1B, 2B, 3A, 3B, 4A, 5B, 5D, 6A, 6B, 6D, 7B, and 7D that were significantly associated with seedling stage resistance. The three SNP markers rs12954, rs34220, and rs42447 on chromosomes 5D, 6A, and 7D, respectively, associated with resistance to *Pt* race PKTTS expressing potential new loci for leaf rust resistance. Overall, this research gives an integrated perspective of leaf rust resistance resources in Iranian bread wheat and recognizes new resistance loci that will be valuable to expand the set of resistance genes available to control this serious disease.

## Introduction

Common wheat (*Triticum aestivum* L.) is among the most important and widely consumed food crops worldwide, and one of the most traded commodities on global markets (FAO 2020). Wheat is frequently attacked by a variety of diseases. Leaf rust caused by *Puccinia triticina* Eriks. (*Pt*), the most prevalent and serious foliar disease impacting wheat production globally, is one of the diseases that causes considerable yield losses in bread wheat (Kolmer 2019; Dinh et al. 2020). In highly susceptible cultivars, the leaf rust fungus mostly affects the leaf blades, but it can also attack the leaf sheath and glumes. Yield loss is usually caused by the reduction of kernel weight and kernel number per spike (Huerta-Espino et al. 2011; Figueroa et al. 2018).

Although fungicides are effective to control rust diseases, using resistant cultivars is more effective, cost-effective, and environmentally safe (Chen, 2020). As a result, having adequate information on the leaf rust agent’s population genetics and identifying novel sources of resistance in the cultivated and landrace gene pools of wheat to contribute to expanding and sustaining the genetic base of leaf rust resistance is critical (McInosch et al. 2013). Plant disease resistance genes can be categorized into two types: all-stage resistance (seedling resistance) and adult-plant resistance (APR). Seedling resistance, which is often race-specific, expresses at all stages of plant development and is commonly associated with a strong hypersensitive reaction with a high level of resistance, despite being easily broken down by changes in rust pathogen virulence. On the other hand, APR that also known as race nonspecific resistance is more effective at adult stages of plant development and is effective against all *Pt* races, and is durable. Many wheat cultivars have become susceptible because of the continual emergence of new pathogen races with new virulence. As a result, new sources of resistance and new *Leaf rust* (*Lr*) resistance genes must be discovered to manage this significant wheat disease (Kolmer et al. 2013; Dinh et al. 2020). Until today a total of 80 *Lr* genes (Leaf Rust Gene) have been discovered (Qureshi et al. 2018; McIntosh et al. 2013; Kumar et al. 2021). The majority of these genes confer seedling resistance, however, nine slow-rusting genes, namely *Lr34* (Dyck 1977), *Lr46* (Singh et al. 1998), *Lr67* (Herrera-Foessel et al. 2014), *Lr68* (Herrera-Foessel et al. 2012), *Lr74* (McIntosh et al. 2013), *Lr75* (Singla et al. 2017), *Lr77* (Kolmer et al. 2018), *Lr78* (Kolmer et al. 2018), and *Lr79* (Qureshi et al. 2018) govern adult plant resistance.

Although bi-parental mapping was successful to discover genomic loci for leaf rust resistance, the restricted recombination events in bi-parental mapping limited the discovery of closely related markers valuable for MAS because of the long linkage block (Riedelsheimer et al. 2012). The genome-wide association study (GWAS) is the most recent methodological technique, which relies on the linkage disequilibrium (LD) principle and the utilization of many SNP (Single Nucleotide Polymorphism) markers. GWAS identifies associations between phenotyping and genotyping data in an association mapping population, and it provides complete surveys of germplasm pools and is a valuable complement to bi-parental mapping research (Zargar et al. 2015; Tibbs Cortes et al. 2021). GWAS utilizes the recombination events that happen during the evolution of populations. This provides the breakup of the LD blocks within the genome and results in a faster decay of the LD in the association mapping than in RILs (recombinant inbred lines) and DH (double haploid) populations, in which only the allelic diversity that separates between the parents can be evaluated. Therefore, GWAS can distinguish associated loci with the trait response at a much higher mapping resolution than bi-parental mapping (Rafalski 2002; Nordborg and Weigel 2008; Zhao et al. 2008; Neumann et al. 2011).

The GWAS method has been successfully applied in different plants for various traits. Different wheat traits have been studied using GWAS including agronomic traits (Safdar et al. 2020; Pang et al. 2020), quality (Yang et al. 2020; Muqaddasi et al. 2020), drought stress (Abou-Elwafa et al. 2021; Shokat et al. 2020; Rahimi et al. 2019), leaf rust (Spakota et al. 2019; Muqaddasi et al. 2021), and stem rust resistance (Saremi et al. 2021; Gao et al. 2017). For leaf rust resistance, Spakota et al. (2019) employed GWAS to identify related genomic areas in wheat genotypes, and eleven QTLs (Quantitative Trait Loci) were identified on nine chromosomes. In wheat landraces, Kertho et al. (2015) observed 73 QTLs associated with resistance to leaf rust and strip rust, and 11 of them were regarded as novel. Also, Gao et al. (2016) discovered 46 QTLs associated with seedling and adult stage resistance for resistance to leaf rust, and about 30% of the phenotypic variance was explained by the ten most significant QTLs.

In the present study, GWAS was conducted on a diverse panel of wheat cultivars and landraces originating from several geographical areas in Iran. This study was designed to detect genetic loci related to seedling resistance to leaf rust by use of 320 Iranian wheat accessions against five *Pt* races, which will be used in marker-assisted selection and further genetic dissections.

## Materials and methods

### Plant materials and *Pt* races

A leaf rust association mapping (AM) panel of 320 wheat accessions was used in the present study, which includes 102 varieties released between 1942 and 2014 and 218 landraces collected between 1931 and 1968 (Supplementary Table 1), along with the susceptible cultivar Boolani. Commercial cultivars were received from the Seed and Plant Improvement Institute (SPII), Karaj, Alborz, Iran, and landraces from the University of Tehran’s Gene Bank. For 298 accessions, both phenotypic and genotypic data were available (90 varieties and 208 landraces).

The five *Pt* races PKTTS, PKTTT, PFTTT, PDTRR, and PDKTT, representing prevalent races of *Pt* in IRAN, were used to screen the wheat accessions. All isolates were collected from bread wheat germplasm. The virulence/avirulence profile of the rust races was determined using infection types based on the seedling stage of Thatcher wheat differentials that are near-isogenic for single-resistance genes based on the race nomenclature of Long and Kolmer (1989). The characteristics of used races are presented in Table 1.

**Table 1.** Virulence/avirulence profile the five *Pt* races used to evaluate the wheat genotypes

| No | Race | Location | Ineffective genes | Effective genes |
| --- | --- | --- | --- | --- |
| 1 | PKTTT | Dezfoul_Khouzestan | *Lr22b, Lr1, Lr2c, Lr3, Lr3ka, Lr3bg, Lr10, Lr11, Lr12, Lr13, Lr14a, Lr14b, Lr15, Lr16, Lr17, Lr18, Lr20, Lr21, Lr22a, Lr23, Lr24, Lr25, Lr26, Lr10, Lr27+ Lr31, Lr28, Lr30, Lr32, Lr33, Lr34, Lr35, Lr36, Lr37, Lrb, Lr13* | *Lr2a, Lr2b, Lr9, Lr19, Lr29* |
| 2 | PFTTT | Dezfoul_Khouzestan | *Lr22b, Lr1, Lr2b, Lr2c, Lr3, Lr3ka, Lr3bg, Lr10, Lr11, Lr12, Lr13, Lr14a, Lr14b, Lr17, Lr18, Lr20, Lr21, Lr22a, Lr23, Lr24, Lr25, Lr26, Lr28, Lr30, Lr32, Lr33, Lr34, Lr35, Lr36, Lr37, Lrb, Lr13* | *Lr2a, Lr9, Lr15, Lr16, Lr19, Lr10, Lr27+ Lr31, Lr29* |
| 3 | PKTTS | Moghan_Ardabil | *Lr22b, Lr1, Lr2c, Lr3, Lr3ka, Lr3bg, Lr10, Lr11, Lr12, Lr13, Lr14a, Lr14b, Lr15, Lr16, Lr17, Lr18, Lr20, Lr21, Lr22a, Lr23, Lr24, Lr25, Lr26, Lr10 / Lr27 + / Lr31, Lr29, Lr30, Lr32, Lr33, Lr34, Lr35, Lr36, Lr37, Lrb, Lr13* | *Lr2a, Lr2b, Lr9, Lr19, Lr28* |
| 4 | PDKTT | Ahwaz_Khouzestan | *Lr22b, Lr1, Lr2c, Lr3, Lr3bg, Lr10, (Lr10, Lr27+Lr31), Lr11, Lr12, Lr13, Lr14a, Lr14b, Lr15, Lr16, Lr17, Lr18, Lr20, Lr21, Lr22a, Lr23, Lr24, Lr25, Lr28, Lr30, Lr32, Lr33, Lr34, Lr35, Lr36, Lr37, Lrb* | *Lr2a, Lr2b, Lr3ka, Lr9, Lr16, Lr19, Lr26, (Lr10, Lr27+Lr10), Lr29* |
| 5 | PDTRR | Gorgan_Golestan | *Lr22b, Lr1, Lr2a, Lr2b, Lr3ka, Lr9, Lr10, Lr11, Lr14a, Lr16, Lr19, Lr20, Lr23, Lr26, (Lr10, Lr27+ Lr31), Lr28, Lr29, Lr33, Lr37, Lr13* | *Lr22b, Lr2c, Lr3, Lr3bg, Lr12, Lr13, Lr14b, Lr15, Lr17, Lr18, Lr21, Lr22a, Lr24, Lr25, Lr30, Lr32, Lr34, Lr35, Lr36, Lrb* |

### Phenotyping at Seedling Stage

Seven seeds of each accession were sown in pots with a diameter and a height of 10 cm, filled with a mixture of common soil, peat moss, and leaf mold. In each pot, four wheat accessions have been positioned at a suitable distance. Then they were stored on a growth chamber at 22-25 °C and a 16 h photoperiod for development. After 8-10 days, when secondary leaves have emerged, inoculation of the seedlings were done separately by the spores of five rust races gathered from various fields of Iran. Then the inoculated seedlings moved in a dark room for one day at 17±2 °C and near 95% moisture, then they were placed in a growth chamber kept at 18°C/20°C (night/day) with 16-h of photoperiod. The 10-12 days after inoculation, plant infection type (IT) was determined based on the method described by McIntosh et al. (1995) rated a scale of 0-4 where 0 = no visible uredia (immune), ; = hypersensitive fleck (very resistant), 1 = small uredia with necrosis (resistant), 2 = small- to medium-sized uredia (resistant to moderately resistant), 3 = medium-sized uredia with or without chlorosis (moderately resistant/moderately susceptible), and 4 = large-sized uredia without chlorosis (susceptible reaction). The 0-4 scale for leaf rust was transformed to a linearized 0-9 scale utilizing the weighted mean of the most and least predominant IT on the same leaf surface to employ the modified McIntosh ITs in genome-wide association studies (GWAS) (Zhang et al. 2014). Values 0 to 6 were considered as resistance IT and, 7 to 9 were considered as susceptible IT.

### Genotyping by sequencing and imputation method

Genotypic evaluation of wheat accessions was conducted in collaboration with the US Ministry of Agriculture and the University of Kansas (Alipour et al. 2017). In brief, genomic DNA of wheat accessions was isolated from young leaves using the modified cetyltrimethyl ammonium bromide (CTAB) method (Saghai-Maroof et al. 1984). The GBS (Genotyping by sequencing) libraries were constructed with two restriction enzymes *Pst*I and *Msp*I according to the method of Poland et al. (2012). Subsequently, barcoded adapters ligation to individual samples were performed using T4 ligase. The DNA purification was carried out using the QIAquick PCR Purification Kit (Qiagen, Inc., Valencia, CA, USA). Finally, the amplified fragments between 250-300 bp were specified on the E-gel system and sent for sequencing on an Ion Proton sequencer (Life Technologies, Inc.). The sequencing data were first trimmed to 64 bp, and the same reads were grouped into tags. The UNEAK GBS pipeline (Lu et al. 2013) as part of the TASSEL 4.0 bioinformatics package (Bradbury et al. 2007) was used for SNPs calling, where SNPs with heterozygosity 10%>, minor allele frequency (MAF) >0.1, and missing data 20%> were removed and other SNPs were used for further analysis. The data was also subjected to imputation using BEAGLE v3.3.2 (Browning and Browning., 2009) based on available allele frequencies obtained after specifying the haplotype phase for all individuals. Four different reference genomes were evaluated and among them, the W7984 reference genome was selected to have the greatest annotation accuracy.

### Phenotypic data analysis

Phenotypic data analysis including descriptive analysis, ANOVA (Analysis of Variance), correlation analysis, and heritability estimation was performed using the SAS software v.9.4. The Shapiro-Wilk test (PROC UNIVARIATE) and Levene’s test (Snedecor and Cochran 1989) were conducted to determine the normal distribution of phenotypic data and to verify the homogeneity of data between experiments, respectively. For the GWAS analysis, the overall mean was used if the data were homogenous. The genetic, environmental, and phenotypic variances were estimated based on the Comstock & Robinson (1952) method as follow:

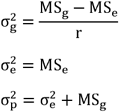

Where MS_g_ is genotype mean square, MS_e_ is error mean square and r is the number of experimental repetitions. The broad-sense heritability for leaf rust was calculated via the ratio of genetic variance to phenotypic variance as follow:

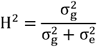

Pearson correlation coefficients among races were determined for IT values based on PROC CORR procedure in SAS software.

### Population structure and LD

To apprehend the genetic structure of the population of Iranian wheat genotypes and to recognize subpopulations, we used Bayesian methods using STRUCTURE v2.3.3 (Pritchard et al. 2000). A putative range of subpopulations starting from k = 1 to 10 was assessed using an admixture model and with a burn-in and simulation phase consisting of 30,000 steps. An adhoc statistic based on the rate of change of the log-likelihood of the data between successive values was used to estimate K. (Evanno et al. 2005; Quraishi et al. 2011). LD between markers was estimated by comparing of observed vs. expected allele frequencies of the markers in TASSEL v.5.2.65 (Bradbury et al. 2007). A Kinship matrix (Q matrix) among individual genotypes for association studies was estimated using all SNP markers; the heat map was performed with the use of a classical equation from Van Randen (2008) in the R software. Principal Component Analysis (PCA) was done by use of SNP markers to specify the genetic relationships between the genotypes, and PC1 was plotted against PC2.

### Genome- wide association mapping

A dataset including 298 accessions was obtained after combining phenotypic (320) and genotypic data (298). GWAS to discover marker-trait associations (MTAs) significantly with seedling resistance was performed using general linear model (GLM) and mixed linear model (MLM) using TASSELv.5.2.65 (Bradbury et al. 2007) and GAPIT package (Lipka et al. 2012) in RStudio (Team 2015). Mixed-linear models (MLM), with kinship matrix (K) and population structure (Q) as a covariate, were selected based on the lowest MSD value. The results using t-tests showed that the GAPIT package (Lipka et al. 2012) supplied stronger control confounding effects. Therefore, only GAPIT results were reported (Lipka et al. 2012). MTAs with a LOD (Logarithm of the Odds) score above 3 (p-value < 0.001) were selected as significant markers for leaf rust resistance. FDR (False Discovery Rate) at the alpha level of 0.05 was used to reduce the false discovery rate of significant markers. In order to reduce the false discovery rate of significant markers, the FDR (False Discovery Rate) was set as 0.05 at the alpha level.

### Gene annotation

The flanking sequences of significant marker-trait associations (MTAs) were received from the Illumina 90K SNP datasets (Wang et al. 2014). Gene ontology (GO) of the sequences significant loci was conducted by use of Ensemble plants database (https://plants.ensembl.org/) by aligning them to the IWGSC RefSeq v1.0 annotation (https://plants.ensembl.org/Multi/Tools/Blast#). The function of the putative genes was determined by examining the metabolic pathways involving the encoded enzymes. The overlapping genes with the highest identity percentage and blast score were selected for further analysis. The information of each gene adjacent to *T. aestivum*, including molecular function, biological process as well as orthologous genes in related species, were obtained from the ensemble-plants database (https://plants.ensembl.org/).

### Comparison QTLs with previously detected *Lr*-gene/QTLs

To discover the relationship between the SNP markers identified that related to leaf rust resistance in this study to previously detected *Lr*-gene/QTLs, the positions of the most significant markers (FDR < 0.05) representative of each QTL to previously mapped QTL/genes were compared using wheat consensus map (Maccaferri et al. 2015). The graphical display of the genetic map was constructed using MapChart (Voorrips 2002).

## Results

### Phenotypic evaluation

IT response (Infection Type) against five pathotypes (PKTTS, PKTTT, PFTTT, PDKTT, and PDTRR) was evaluated in the greenhouse for 320 accessions. The results are presented in Supplementary Table 2. In all the experiments, the susceptible cultivar Boolani was highly infected and showed the expected compatible ITs of 3 to 4 for all five pathotypes. Wheat accessions had a wide variety of responses to all five *Pt* races used in our research (Supplementary Table 2). The leaf rust scores varied from immune (IT=0, LS=0) to highly susceptible reaction (IT=4, LS=9) to all five *Pt* races (Table 2). The majority of the tested wheat accessions were susceptible to the *Pt* races and of these, 36, 32, 59, 38, and 77 accessions were resistant (IT rating < 3, linear score < 8) to races PKTTS, PKTTT, PFTTT, PDKTT, and PDTRR, respectively (Table 2), also a total of ten accessions were resistant to all five pathotypes (Table 3).

**Table 2.**
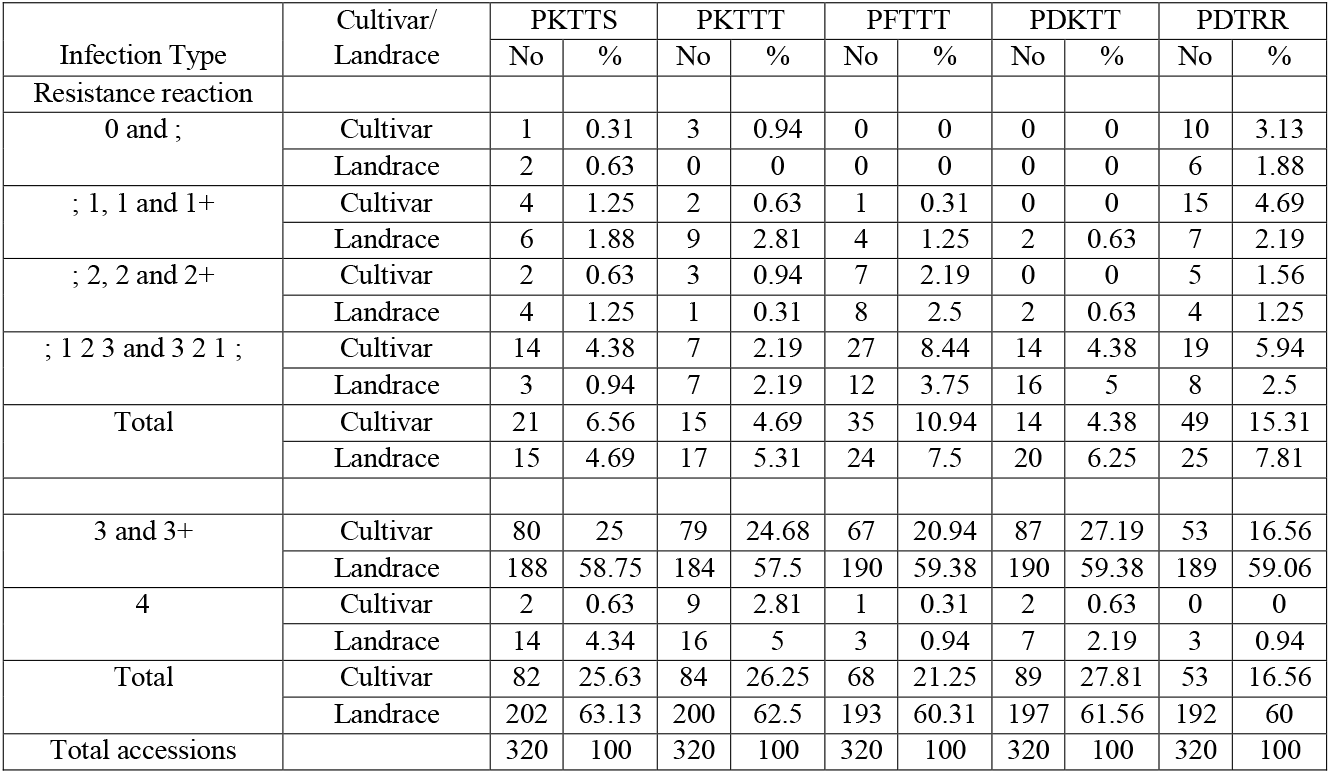
grouping of wheat population based on infection type to five Pt races

**Table 3.** Resistance wheat accessions to all five *Puccinia triticina* (*Pt*) races

| Accession | Origin | Type | Disease score |  |  |  |  |
| --- | --- | --- | --- | --- | --- | --- | --- |
|  |  |  | PKTTT | PFTTT | PDKTT | PKTTS | PDTRR |
| 622084 | Mazandaran_Sari | Landrace | 0.70 | 0.70 | 5.33 | 0.70 | 0.70 |
| 622099 | Gilan_Rasht | Landrace | 0.70 | 3.34 | 2.67 | 0.70 | 0.35 |
| 622247 | Mazandaran_Sari | Landrace | 0.70 | 0.70 | 5.33 | 0.35 | 0.67 |
| 622264 | Mazandaran_Babol | Landrace | 0.70 | 2.17 | 1.19 | 1.67 | 0.35 |
| 622272 | Mazandaran_Amol | Landrace | 1.19 | 4.00 | 2.17 | 0.35 | 0.35 |
| 624381 | Bakhtaran_Bakhtaran | Landrace | 0.70 | 1.67 | 4.00 | 0.70 | 0.35 |
| 627856 | Mazandaran_Sari | Landrace | 1.19 | 0.70 | 1.19 | 1.19 | 0.35 |
| 627963 | Hamedan_Hamedan | Landrace | 0.70 | 1.19 | 1.69 | 1.50 | 0.35 |
| 627057 | Gilan_Fooman | Landrace | 2.17 | 1.84 | 7.5 | 2.84 | 0 |
| Shinghai | - | Varity | 6.00 | 4.00 | 7.67 | 7.67 | 0.35 |

The results of the Shapiro-Wilk normality test indicated that the phenotypic data of all five *Pt* pathotypes deviated significantly from a normal distribution (Table 4). The Leven’s test was then performed to test the homogeneity of the data. The results of Levene’s test indicated that the phenotypic variance of the data within experiments was homogenous (P = 0.23 to 0.79) for all five *Pt* pathotypes (Table 4). Therefore, the overall mean for each wheat accession was calculated and utilized in GWAS study.

**Table 4.** Descriptive statistics of 320 wheat accessions evaluated for their response to five *Puccinia triticina* (*Pt*) races

| Race | Mean | Min | Max | SD | Shapiro-Wilk test<sup>a</sup> | Leven's test<sup>b</sup> | $\sigma_g^2$ | $\sigma_e^2$ | $\sigma_p^2$ | H<sup>2</sup> (%) |
| --- | --- | --- | --- | --- | --- | --- | --- | --- | --- | --- |
| PKTTT | 8.30 | 0 | 9.00 | 2.15 | P<0.0001 | P=0.595 | 4.604 | 0.017 | 4.621 | 99.63 |
| PFTTT | 7.92 | 0.70 | 9.00 | 2.00 | P<0.0001 | P=0.229 | 3.835 | 0.394 | 4.23 | 90.67 |
| PKTTS | 8.30 | 0 | 9.00 | 2.06 | P<0.0001 | P=0.792 | 4.173 | 0.115 | 4.29 | 97.34 |
| PDTRR | 7.33 | 0 | 9.00 | 3.12 | P<0.0001 | P=0.667 | 9.62 | 0.167 | 9.79 | 99.48 |
| PDKTT | 8.37 | 1.19 | 9.00 | 1.11 | P<0.0001 | P=0.779 | 0.217 | 1.13 | 1.347 | 83.89 |

<sup>a</sup> Shapiro-Wilk test was conducted to determine if the phenotypic data were normal or not. p<0.05 shows non-normal distribution.
<sup>b</sup> Leven's test was conducted to determine if the data among the experiments are homogenous or not. P>0.05 shows equal variance.
SD = standard deviation
$\sigma_g^2$ =estimates of genotypic variance
$\sigma_e^2$ =estimates of environmental variance
$\sigma_p^2$ =estimates of phenotypic variance
H<sup>2</sup> = broad-sense heritability

The ANOVA for leaf rust seedling reactions showed highly significant differences (P < 0.001) between races, accessions, and race × accession interaction (Table 5). The coefficient of correlation (r) among all five *Pt* pathotypes was highly positive and significant. The correlation coefficient values for ITs ranged from 0.40-0.71. In particular, high correlation coefficient values were observed for the pair-correlations of PKTTS vs. PFTTT (0.71), PFTTT vs. PDTRR (0.69), PFTTT vs. PKTTT (0.65), and PKTTS vs. PDTRR (0.60) (Table 6).

**Table 5.**
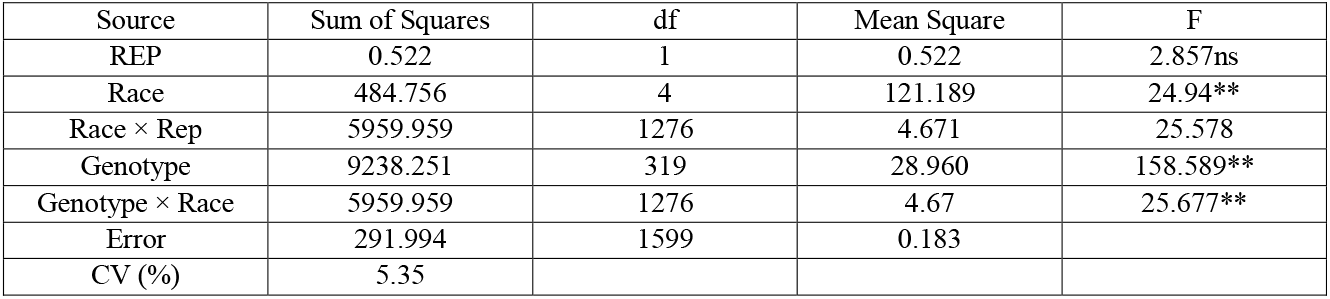
Combined analysis of variance for infection types of wheat accessions to five *Pt* races

| Source | Sum of Squares | df | Mean Square | F |
| --- | --- | --- | --- | --- |
| REP | 0.522 | 1 | 0.522 | 2.857ns |
| Race | 484.756 | 4 | 121.189 | 24.94** |
| Race × Rep | 5959.959 | 1276 | 4.671 | 25.578 |
| Genotype | 9238.251 | 319 | 28.960 | 158.589** |
| Genotype × Race | 5959.959 | 1276 | 4.67 | 25.677** |
| Error | 291.994 | 1599 | 0.183 |  |
| CV (%) | 5.35 |  |  |  |

**Table 6.**
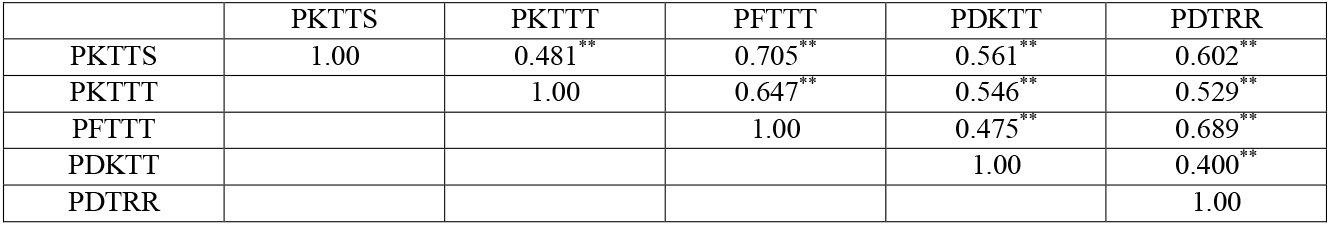
Correlation coefficients between the phenotypic data of 320 wheat accessions evaluated for response to *Puccinia triticina* (*Pt*) races

|  | PKTTS | PKTTT | PFTTT | PDKTT | PDTRR |
| --- | --- | --- | --- | --- | --- |
| PKTTS | 1.00 | 0.481** | 0.705** | 0.561** | 0.602** |
| PKTTT |  | 1.00 | 0.647** | 0.546** | 0.529** |
| PFTTT |  |  | 1.00 | 0.475** | 0.689** |
| PDKTT |  |  |  | 1.00 | 0.400** |
| PDTRR |  |  |  |  | 1.00 |

### Linkage disequilibrium

Linkage disequilibrium decay was examined for the original and imputed datasets for three genomes separately and all chromosomes within each genome. Based on the linkage disequilibrium analysis, the LD declined with the increases in genetic distance. The significant marker pairs at P < 0.001 were considered for the study. In general, genome B and D had the highest and lowest marker density, respectively (Table 7 and 8). However, it is more useful to test the LD between each pair of SNPs located on the same chromosome and determine the average of the LD in each genome to identify the pattern of LD in the three genomes. At the genome level in original datasets, for both Landraces and varieties, Genome A had 22.34% of significant marker pairs with an average r^2^*-* value of 0.10 for varieties and 31.58% of significant marker pairs with an average r^2^*-* value of 0.1 for landraces. The maximum marker density for both Landraces and varieties was observed on chromosome 2B with 31387 pair SNPs for varieties and 30754 pair SNPs for landraces. Genome B had 26.39% of significant markers with an average r^2^*-* value of 0.13 for varieties and 28.71% of significant markers with an average r^2^*-* value of 0.078 for landraces. Genome D had 24.34% of significant marker pairs with an average r^2^*-* value of 0.12 for varieties and 25.27% of significant marker pairs with an average r^2^*-* value of 0.1 for landraces.

**Table 7.** A summary of observed LD (r^2^) among SNP pairs and the number of significant SNP pairs per chromosomes and genomes of Iranian bread wheat cultivars and landraces in original datasets

| Chromosome | Cultivar |  |  |  | Landrace |  |  |  |
| --- | --- | --- | --- | --- | --- | --- | --- | --- |
| | TNSP | $r^2$ | Distance (cM) | NSSP | TNSP | $r^2$ | Distance (cM) | NSSP |
| 1A | 16283 | 6.75189 | 0.109825 | 3917 (24.06%) | 12992 | 11.528 | 0.114653 | 4750 (36.53%) |
| 1B | 22004 | 4.47251 | 0.141322 | 5775 (6.25%) | 25210 | 4.5844 | 0.089915 | 8237 (32.67%) |
| 1D | 10733 | 10.2787 | 0.184493 | 3367 (31.37%) | 19042 | 6.3896 | 0.071123 | 4235 (22.24%) |
| 2A | 20435 | 5.05017 | 0.123509 | 4915 (24.05%) | 22359 | 4.8757 | 0.114976 | 7734 (34.59%) |
| 2B | 31387 | 4.11067 | 0.12831 | 8386 (26.72%) | 30754 | 4.125 | 0.092202 | 9932 (32.29%) |
| 2D | 13331 | 6.58741 | 0.266553 | 4494 (33.7%) | 15780 | 6.7303 | 0.195837 | 5174 (32.78%) |
| 3A | 17793 | 9.80947 | 0.098266 | 3567 (20.05%) | 17858 | 9.1444 | 0.069792 | 4272 (23.92%) |
| 3B | 28610 | 4.62157 | 0.129764 | 7815 (27.32%) | 29925 | 4.4702 | 0.091393 | 9351 (31.25%) |
| 3D | 4725 | 17.5873 | 0.097313 | 704 (14.90%) | 7601 | 19.707 | 0.090504 | 1628 (21.42%) |
| 4A | 15937 | 7.97345 | 0.130939 | 3621 (22.72%) | 15490 | 8.2723 | 0.109944 | 4342 (28.03%) |
| 4B | 8325 | 9.53805 | 0.100295 | 1820 (21.86%) | 8450 | 10.408 | 0.050408 | 1373 (16.25%) |
| 4D | 3001 | 25.9259 | 0.134246 | 558 (18.59%) | 3171 | 25.715 | 0.10865 | 1088 (34.31%) |
| 5A | 15117 | 7.76999 | 0.110108 | 3238 (21.42%) | 16814 | 8.0587 | 0.080919 | 4825 (28.70%) |
| 5B | 26207 | 6.39664 | 0.132941 | 7680 (29.31%) | 26766 | 6.3968 | 0.069826 | 6575 (24.56%) |
| 5D | 5946 | 25.1631 | 0.00939E | 796 (13.39%) | 6990 | 28.565 | 0.066032 | 1377 (19.70%) |
| 6A | 16119 | 7.47835 | 0.106207 | 3117 (19.34%) | 17460 | 7.3983 | 0.118612 | 6784 (38.80%) |
| 6B | 21869 | 4.38736 | 0.139339 | 6339 (28.99%) | 24908 | 4.6098 | 0.071623 | 6560 (26.64%) |
| 6D | 7845 | 18.2414 | 0.0104 | 1358 (17.31%) | 8963 | 17.787 | 0.078422 | 2160 (24.10%) |
| 7A | 22236 | 6.02434 | 0.015 | 5304 (23.85%) | 27109 | 6.1406 | 0.107505 | 8369 (30.87%) |
| 7B | 24351 | 4.78852 | 0.110689 | 5149 (21.14%) | 25094 | 4.6028 | 0.080258 | 7081 (28.22%) |
| 7D | 8108 | 19.9858 | 0.0166 | 1794 (22.13%) | 10344 | 19.537 | 0.094395 | 2508 (24.24%) |
| A genome | 123920 | 7.26539 | 0.09912 | 27679 (22.3%) | 130082 | 7.9166 | 0.102343 | 41076 (31.6%) |
| B genome | 162753 | 5.47362 | 0.12609 | 42964 (26.4%) | 171107 | 5.5997 | 0.07795 | 49109 (28.7%) |
| D genome | 53689 | 17.6814 | 0.11827 | 13071 (24.4%) | 71891 | 17.776 | 0.10071 | 18170 (25.3%) |
| Total | 340362 | 10.1401 | 0.11449 | 83714 (25 %) | 373080 | 10.430 | 0.09367 | 108355 (29 %) |

TNSP: Total number of SNP pairs, NSSP: Number of significant SNP pairs (P value< 0.001)

**Table 8.** A summary of observed LD (r^2^) among SNP pairs and the number of significant SNP pairs per chromosomes and genomes of Iranian bread wheat cultivars and landraces in imputed datasets

| Chromosome | Cultivar |  |  |  | Landrace |  |  |  |
| --- | --- | --- | --- | --- | --- | --- | --- | --- |
| | TNSP | $r^2$ | Distance (cM) | NSSP | TNSP | $r^2$ | Distance (cM) | NSSP |
| 1A | 85575 | 0.148218 | 1.737691 | 27125 (31.7%) | 92925 | 0.112764 | 1.596397 | 33515 (36.07%) |
| 2A | 118025 | 0.292156 | 0.974187 | 57858 (49.02%) | 123175 | 0.297454 | 0.944378 | 68675 (55.75%) |
| 3A | 83675 | 0.159365 | 2.576447 | 25903 (30.96%) | 73525 | 0.136413 | 2.939734 | 28144 (38.28%) |
| 4A | 114925 | 0.371766 | 1.513597 | 57774 (50.27%) | 108375 | 0.376224 | 1.612148 | 65451 (60.39%) |
| 5A | 59375 | 0.169369 | 2.383461 | 18718 (31.53%) | 58475 | 0.150278 | 2.416511 | 24007 (41.06%) |
| 6A | 85175 | 0.181387 | 1.487802 | 29645 (34.8%) | 84425 | 0.181735 | 1.501019 | 40176 (47.59%) |
| 7A | 128575 | 0.234215 | 1.344495 | 49426 (38.44%) | 126575 | 0.214252 | 1.365959 | 63357 (50.05%) |
| 1B | 131075 | 0.206251 | 1.063813 | 49717 (37.93%) | 133525 | 0.157517 | 1.041252 | 63803 (47.78%) |
| 2B | 165475 | 0.198105 | 0.859164 | 66129 (39.96%) | 155625 | 0.177663 | 0.913543 | 78536 (50.46%) |
| 3B | 176175 | 0.245726 | 0.876581 | 78363 (44.48%) | 170925 | 0.221549 | 0.903978 | 89150 (52.16%) |
| 4B | 51325 | 0.1455 | 2.516753 | 13477 (26.26%) | 43025 | 0.1018 | 3.002768 | 12311 (28.61%) |
| 5B | 134225 | 0.204683 | 1.433217 | 55633 (41.45%) | 134675 | 0.14301 | 1.449279 | 56285 (41.79%) |
| 6B | 158275 | 0.205457 | 0.788418 | 66108 (41.77%) | 164475 | 0.139023 | 0.758663 | 71582 (43.52%) |
| 7B | 132875 | 0.156677 | 1.102364 | 41160 (30.98%) | 125875 | 0.129711 | 1.157535 | 50573 (40.18%) |
| 1D | 37075 | 0.294821 | 4.409069 | 16539 (44.61%) | 40975 | 0.232567 | 3.832101 | 19755 (48.21%) |
| 2D | 48025 | 0.23446 | 2.2455 | 16275 (33.89%) | 52825 | 0.169092 | 2.048568 | 20548 (38.9%) |
| 3D | 25475 | 0.143085 | 6.286093 | 5413 (21.25%) | 30125 | 0.174879 | 5.31564 | 11411 (37.88%) |
| 4D | 10275 | 0.167587 | 10.56621 | 2189 (21.3%) | 10375 | 0.14746 | 10.71346 | 3543 (34.15%) |
| 5D | 22375 | 0.155406 | 9.337668 | 5503 (24.59%) | 24825 | 0.142184 | 8.361416 | 8953 (36.06%) |
| 6D | 28475 | 0.142966 | 5.369092 | 6844 (24.04%) | 33475 | 0.14123 | 4.565844 | 12606 (37.66%) |
| 7D | 34475 | 0.208327 | 5.795738 | 10809 (31.35%) | 40475 | 0.153099 | 4.947296 | 14019 (34.64%) |
| A genome | 675325 | 0.235213 | 1.620443 | 266449 (39.4%) | 667475 | 0.223484 | 1.64269 | 323325 (48.4%) |
| B genome | 949425 | 0.20158 | 1.083656 | 370587 (39.0%) | 928125 | 0.160951 | 1.110386 | 422240 (45.5%) |
| D genome | 206175 | 0.205106 | 5.343207 | 63572 (30.83%) | 233075 | 0.170391 | 4.707401 | 90835 (38.97%) |
| Total | 1830925 | 0.214383 | 1.761302 | 700608 (38.3%) | 1828675 | 0.184979 | 1.76314 | 836400 (45.7%) |

TNSP: Total number of SNP pairs, NSSP: Number of significant SNP pairs (P value < 0.001)

In imputed datasets, the extent of LD for the wheat varieties and landraces was 0.21 and 0.18, respectively, and the average genetic distance for both of them was about 1.76 cM. At the chromosome level, the maximum marker density for both Landraces and varieties was observed on chromosome 3B with 176175 pair SNPs for varieties and 170925 pair SNPs for landraces. In general, the proportion of each A, B, and D genomes from total pairwise varieties SNP markers were estimated at almost 39, 39, and 31%, respectively, and in the landraces SNP markers approximately 48, 45, and 40%, respectively.

### Population structure and kinship matrix

In order to determine the appropriate number of subpopulations, the number of clusters was plotted (K) against ΔK. The largest ΔK value was observed at K = 3 suggesting the presence of three subpopulations in the tested accessions for both datasets (Figure 1). Using the structure software, the population of 286 accessions was structured into three subpopulations, Sub1, Sub2, and Sub_3 (Figure 2). Sub_1 included 84 accessions, Sub_2 included 75 accessions and Sub_3 included 127 accessions.

**Fig 1.**
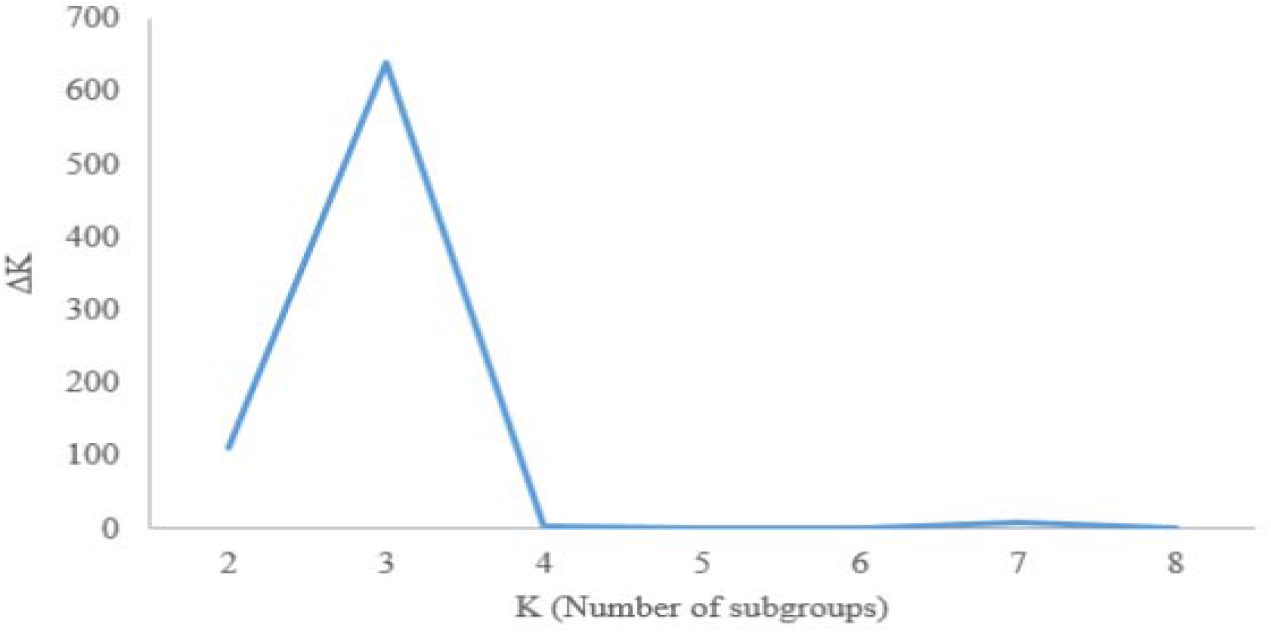
Determination of subpopulations number in wheat genotypes based on ΔK values

**Fig 2.**
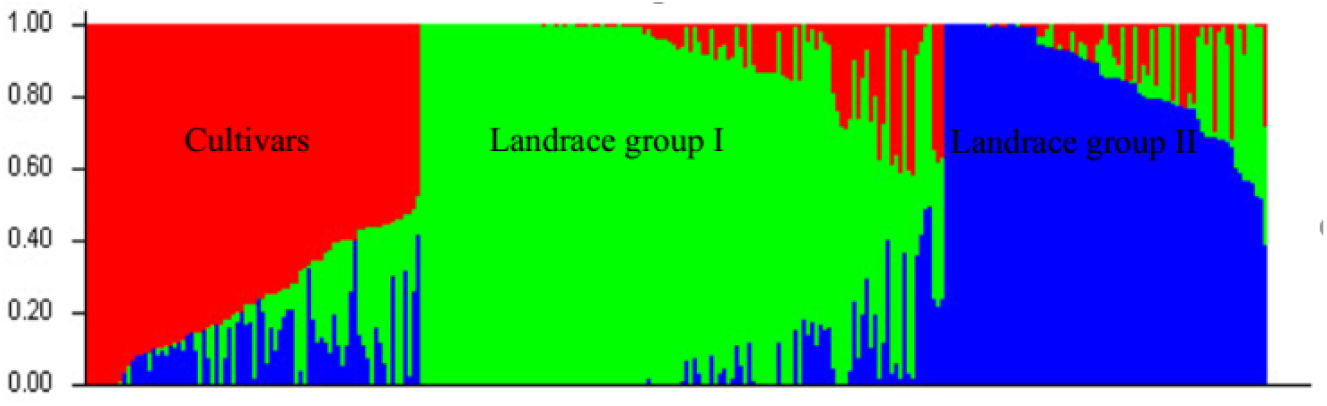
A structure plot of the 286 wheat genotypes and landraces determined by K=3

To better evaluate population structure and investigate genetic relationships among wheat accessions, PCA of original and imputed SNPs was performed in 286 wheat accessions. For the original datasets, the two major components described a total of 18.59% of the genetic variance (Figure 3a), whereas it was 23.1% for the imputed datasets (Figure 3b). Group 1 included 105 accessions with 71 varieties and 34 landraces (63.28%); Group 2 included the 108 accessions with 102 landraces and 6 varieties (37.76%); Group 2 included the smallest number of accessions with 73 accessions with 62 landraces and 11 varieties (25.52%) (Figure 4a). For Original datasets, accessions were also clustered into three main groups. Group 1 included 116 accessions with 6 varieties and 110 landraces; Group 2 included 103 accessions with 85 landraces and 18 varieties; Group 3 included 66 accessions with 3 landraces and 63 varieties (Figure 4b).

**Fig 3.**
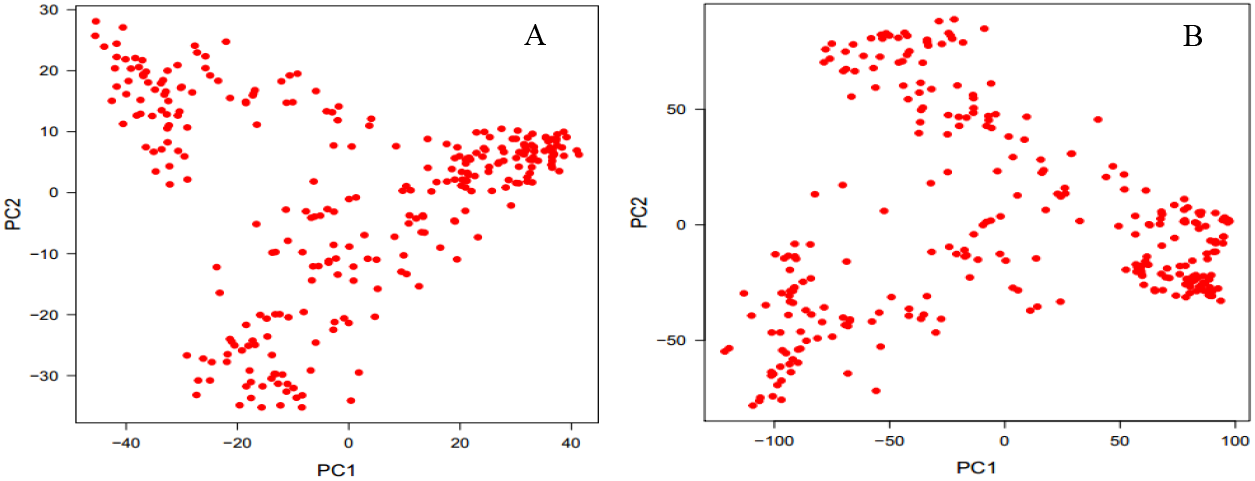
Principal component analysis of Iranian accessions using original SNPs (A), and imputed SNPs.

**Fig 4.**
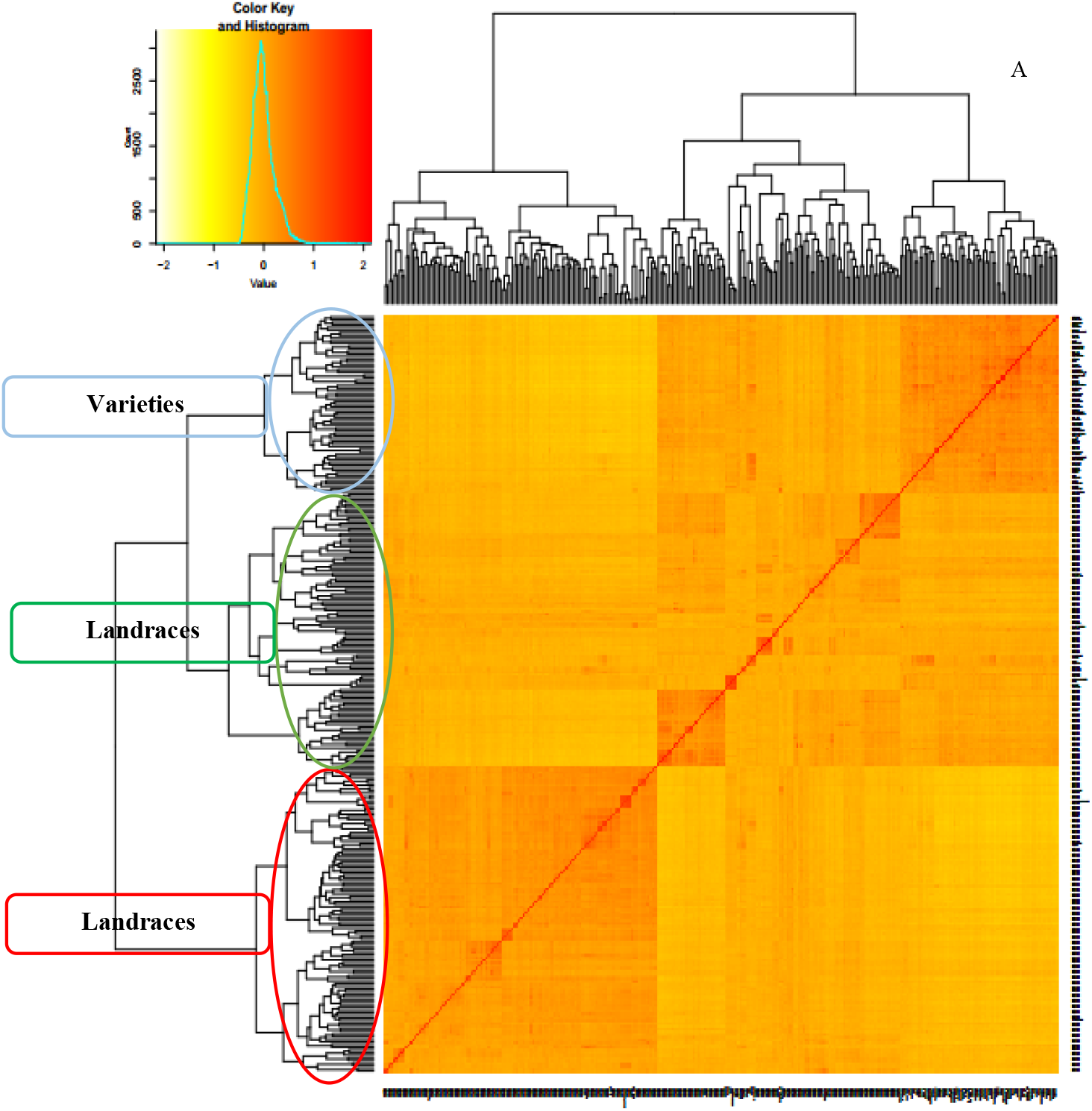
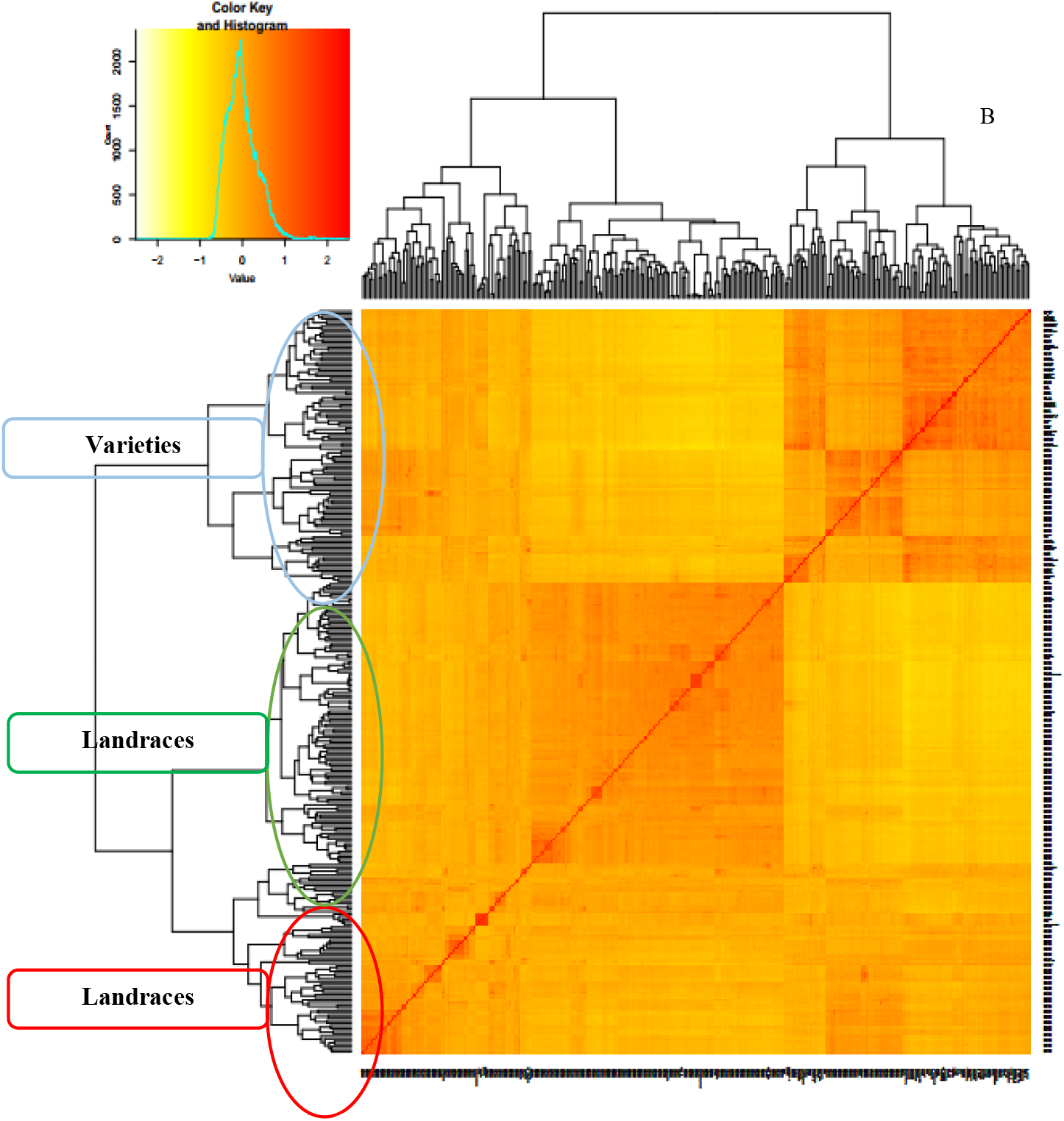
Cluster analysis using kiniship matrix of original data (A) and imputed data (B) for Iranian wheat accessions

### Marker - trait associations

GWAS was conducted using infection type data to reveal the association between the phenotypic and genotypic data in the seedling stage. A total of 9043 and 44106 SNP markers were used in GWAS analysis in original and imputed datasets, respectively. Generally, GWAS identified a total of 36 and 390 significant marker-trait associations for original and imputed datasets at a significance level of –log10 P >3 (P<0.001), respectively (Table 9). In original datasets, 7, 4, 18, 3, and 4 significant SNP were detected for resistance to the pathotypes PKTTS, PKTTT, PFTTT, PDTRR, and PDKTT, respectively (Supplementary Table 3). These SNPs were distributed on 1B, 2A, 2B, 3B, 4A, 4B, 4D, 5B, 5D, 6A, 6D, 7B and 7D chromosomes. In imputed datasets, 137, 101, 48, 45 and, 59 significant SNP were detected for resistance to the races PKTTS, PKTTT, PFTTT, PDTRR, and PDKTT, respectively (Supplementary Table 4). These SNPs were distributed on all chromosomes. In imputed datasets, rs10560, rs12690, rs12954, rs14228, rs14431, rs17878, rs18054, rs19727, rs21735, rs21939, rs22627, rs23335, rs23336, rs23337, rs28088, rs28089, rs28358, rs38875, rs44015, rs44160, rs44883, rs45575, rs47218, rs58203, rs59576, rs61015, rs61600, rs62825, rs6313, rs6314, rs64792, rs7195, rs8909, and rs9493 markers were significant for resistance at least two races, while the remaining MTAs were significant to only a single race (Supplementary Table 4). The largest number of associated markers in both datasets was identified on the B genome whereas the smallest number of significant SNPs markers for original and imputed datasets were on the D genome. The major of MTAs in imputed and original datasets were identified on chromosome 2A (52 MTAs) and 1B (6 MTAs), respectively.

**Table 9.**
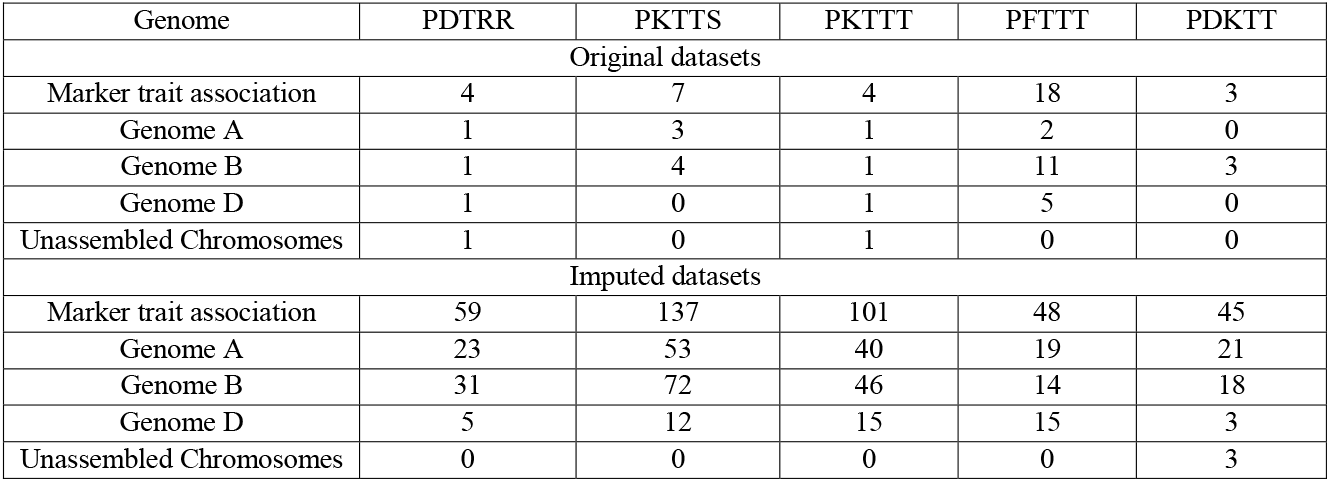
Number of Marker-trait associations (MTAs) for infection type of studied races in Iranian wheat genotypes (P Value < 0.001)

| Genome | PDTRR | PKTTS | PKTTT | PFTTT | PDKTT |
| --- | --- | --- | --- | --- | --- |
| Original datasets |  |  |  |  |  |
| Marker trait association | 4 | 7 | 4 | 18 | 3 |
| Genome A | 1 | 3 | 1 | 2 | 0 |
| Genome B | 1 | 4 | 1 | 11 | 3 |
| Genome D | 1 | 0 | 1 | 5 | 0 |
| Unassembled Chromosomes | 1 | 0 | 1 | 0 | 0 |
| Imputed datasets |  |  |  |  |  |
| Marker trait association | 59 | 137 | 101 | 48 | 45 |
| Genome A | 23 | 53 | 40 | 19 | 21 |
| Genome B | 31 | 72 | 46 | 14 | 18 |
| Genome D | 5 | 12 | 15 | 15 | 3 |
| Unassembled Chromosomes | 0 | 0 | 0 | 0 | 3 |

The results of FDR ≤ 0.05 of the GWAS results of both datasets are shown in Table 10. The results showed that there are only two markers for the original datasets in FDR < 0.05. Two identified markers (rs7087 and rs7088) are associated with the PFTTT race located on chromosome 2B and 6D at 59.184 cM and 51.214 cM, respectively. The results of the imputed datasets showed that there are a total of 17 MTAs in the FDR less than 0.05. All of the MTAs except three MTAs included rs9493, rs62902, and rs62903 (PDKTT), were assigned to PKTTS race. These MTAs were distributed on 1B, 2B, 3A, 3B, 4A, 5B, 5D, 6A, 6B, 6D, 7B and, 7D chromosomes. The maximum of MTAs (4 MTAs) were located on chromosome 1B. The results of Manhattan and QQ-plots of highly associated SNPs for infection type are presented in Figure 5.

**Table 10.** Summary of marker trait associations (MTAs) discovered significant for resistance to *Puccinia triticina* (Pt) races PKTTT, PFTTT, PKTTS, PDKTT, and PDTRR at FDR < 0.05

| SNP | Race | Type data | Allele | Chromosome | Position (cM) | P.value | maf | R<sup>2</sup> | pFDR | effect |
| --- | --- | --- | --- | --- | --- | --- | --- | --- | --- | --- |
| rs7088 | PFTTT | Original | C/T | 2B | 59.184 | 7.53E-07 | 0.23776 | 0.264 | 0.0068 | 0.16 |
| rs7087 | PFTTT | Original | C/T | 6D | 51.214 | 2.97E-06 | 0.21503 | 0.264 | 0.0134 | 0.16 |
| rs43242 | PKTTS | Imputed | G/T | 1B | 30.143 | 3.32E-08 | 0.12237 | 0.348 | 0.0015 | 1.45 |
| rs45675 | PKTTS | Imputed | T/C | 1B | 30.711 | 3.51E-06 | 0.08391 | 0.348 | 0.0296 | 1.29 |
| rs45676 | PKTTS | Imputed | C/T | 1B | 30.711 | 5.20E-06 | 0.08566 | 0.348 | 0.0296 | 1.25 |
| rs43873 | PKTTS | Imputed | A/G | 1B | 45.006 | 6.78E-06 | 0.07867 | 0.348 | 0.0311 | 1.62 |
| rs62679 | PKTTS | Imputed | A/G | 3A | 57.649 | 7.04E-06 | 0.09965 | 0.348 | 0.0311 | 0.94 |
| rs28322 | PKTTS | Imputed | C/T | 3B | 62.576 | 5.37E-06 | 0.19405 | 0.348 | 0.0296 | 1.09 |
| rs3229 | PKTTS | Imputed | C/T | 3B | 62.576 | 1.18E-05 | 0.18881 | 0.348 | 0.0433 | -1.15 |
| rs10560 | PKTTS | Imputed | C/T | 4A | 75.832 | 2.50E-06 | 0.07167 | 0.348 | 0.0296 | 1.39 |
| rs21735 | PKTTS | Imputed | C/G | 4A | 62.152 | 2.92E-06 | 0.08042 | 0.348 | 0.0296 | -1.50 |
| rs6151 | PKTTS | Imputed | A/C | 5B | 70.681 | 4.52E-06 | 0.15559 | 0.348 | 0.0296 | -0.91 |
| rs12954 | PKTTS | Imputed | A/G | 5D | 111.553 | 1.56E-05 | 0.11188 | 0.348 | 0.0492 | 1.00 |
| rs34220 | PKTTS | Imputed | A/C | 6A | 10.247 | 1.55E-05 | 0.06118 | 0.348 | 0.0492 | 1.18 |
| rs15705 | PKTTS | Imputed | A/G | 7B | 117.414 | 8.23E-06 | 0.35664 | 0.348 | 0.0330 | -0.85 |
| rs42447 | PKTTS | Imputed | A/G | 7D | 83.31 | 1.06E-06 | 0.19755 | 0.348 | 0.0234 | 1.01 |
| rs9493 | PDKTT | Imputed | A/G | 3B | 22.764 | 7.73E-07 | 0.09790 | 0.185 | 0.0243 | 0.51 |
| rs62902 | PDKTT | Imputed | C/G | 6B | 47.831 | 1.45E-06 | 0.13286 | 0.185 | 0.0243 | 0.47 |
| rs62903 | PDKTT | Imputed | C/G | 6B | 47.831 | 1.65E-06 | 0.13461 | 0.185 | 0.0243 | 0.47 |

**Fig 5.**
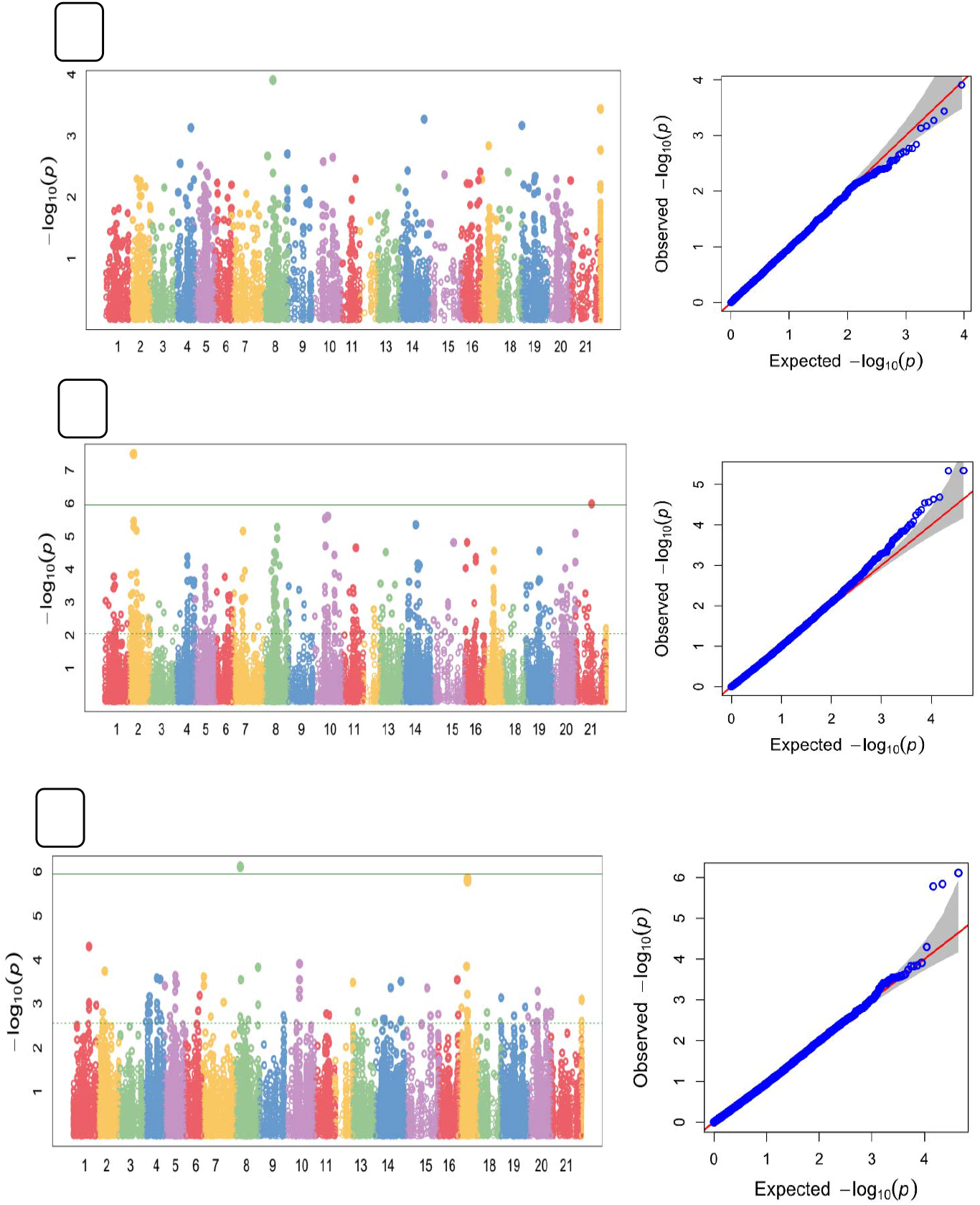
Manhattan and QQ-plots of highly associated haplotypes for Leaf rust. A) PFTTT race, B) PKTTS race, C) PDKTT race. The numbers of 1-22 on X axis represents chromosomes 1A, 1B, 1D, 2A, 2B, 2D, 3A, 3B, 3D, 4A, 4B, 4D, 5A, 5B, 5D, 6A, 6B, 6D, 7A, 7B, 7D, and unknown respectively.

### Gene annotation

To gain a deeper understanding of the relationship between SNPs and leaf rust resistance, we examined the gene annotations of these SNPs and studied the effect of SNPs on genes (Tables 15 and 16). The results of gene ontology showed that of the 390 MTAs that we identified using the imputed datasets 24.62% of them were located within protein-coding genes (Supplementary Table 5). For the original datasets, 6 MTAs (16.67%) were found within protein-coding genes (Supplementary Table 6). The chromosomal sequence, chromosomal position, the closest wheat gene to them, molecular function and biological processes of these genes, and other information of MTAs are presented in Tables 15 and 16. These genes mostly encode proteins involved in nucleotide binding, hydrolase activity, potassium ion transmembrane transporter activity, hydrolase activity, ATP binding, fatty-acyl-CoA binding, lipid binding, hydrolase activity, protein kinase activity, hydrolyzing O-glycosyl compounds, beta-fructofuranosidase activity, acting on glycosyl bonds and, protein binding.

## Discussion

The development of new races of leaf rust pathogens is a constant threat to global wheat production. Therefore, it is necessary to investigate additional resistance sources and genes to generate cultivars with effective genes for resistance to leaf rust. GWAS is a potent strategy to recognize QTL associated with complex traits in plants (Alqudah et al. 2020; Hall et al. 2010). GWAS has been successfully applied in wheat gene pools to identify several genes/QTLs that contribute to leaf rust resistance at both the seedling and adult plant stages (Kertho et al. 2015; Aoun et al. 2016; Turner et al. 2017; Riaz et al. 2018). As shown in the present research and previous studies, wheat landraces are a rich source of genes for resistance to leaf rust (Kertho et al. 2015; Aoun et al. 2016; Turner et al. 2017; Riaz et al. 2018). In the present study, we recognized ten accessions resistant to five *Pt* races that are prevalent in Iran. Nine of these accessions were wheat landraces. Iran is one of the countries in the Fertile Crescent region, which is known as the center origin and diversity of wheat. In addition, previous studies have suggested that the center of origin of P. triticina is probably somewhere in the Fertile Crescent region in southwest Asia (Arthur, 1929), where both sexual and asexual reproduction common (Kolmer et al. 2011). However, possible sexual recombination events are rare in the world (Kolmer et al. 2011). Therefore, this region could provide an opportunity for natural selection and maintenance of resistance accessions. Although wheat landraces may exhibit less desirable agronomic traits, they have been cultivated over many years by local farmers and have been adapted to climate conditions, and have been evolved disease resistance. They also are relatively easy to use in breeding programs compared to alien species (Sehgal et al. 2016). Therefore, the resistant landraces identified in the present study should be useful for developing wheat cultivars resistant to leaf rust.

Pearson correlation coefficients based on infection types revealed the presence of significant correlations for all races in this study (Table 6). These significant correlations were mainly attributed to the similar *Pt* populations across the country and similar virulence/avirulence profile of these races. According to the virulence/avirulence profile test performed on 20 wheat lines carrying a single *Lr* gene, these five races were virulent to *Lr22b, Lr1, Lr3ka, Lr9, Lr10, Lr11, Lr14a, Lr20, Lr23, Lr26, Lr33, Lr37*, and *Lr13* genes. Also, the GWAS panel used in this study probably controls same genomic loci conferring resistance to five *Pt* races, and this was further proved by the GWAS analysis results that permitted identification of common QTL, for example, rs38875 marker, conferring resistance to three *Pt* races PFTTT, PDKTT, and, PDTRR (Supplementary Table 4). Also, the rs59576 marker confers resistance to three *Pt* races PFTTT, PDKTT, and, PKTTS. Research findings by Desidrio et al. (2014) and Sapkota et al. (2019) showed that there is a high correlation between the phenotypic data evaluated with several *Pt* breeds, and common genomic loci were identified for resistance of those breeds, which was consistent with the results of this study.

Information about population structure as a confounding factor plays an important role in GWAS analysis because the presence of population structure in the GWAS panel can lead to false association results (Oraguzie et al. 2007). Selection and Genetic drift are two important factors that justify the presence of a subpopulation in a large population (Buckler and Thornsberry 2002). Population structure, kinship matrix, and PCA analysis are widely utilized approach to infer cryptic population structure from genome-wide data such as high-density SNPs. In the present study, population STRUCTURE, PCA analysis, and kinship matrix classified the wheat accessions into three major subpopulations in both the original and imputed datasets. The population structure recognized in this study had a lesser number of subpopulations than several previously reported GWAS studies (Li et al. 2016; Liu et al. 2017; Zegeye et al. 2014), that this due to all of the accessions obtained a small region. The presence of structure in the current population is for two reasons. A significant number of wheat cultivars in this study were obtained from the International Center for Maize and Wheat Improvement (CIMMYT), which is used either directly or as parents in cross-breeding programs leading to new cultivars (Supplementary Table 1). For both original and imputed datasets, population structure showed that CIMMYT advanced lines like Chamran, Darab 2, and Gahar appeared in the same sub-populations along with Iranian cultivars. Also, the role of agro-ecological zones of the country in the formation of three sub-populations and the preservation of this genetic diversity, especially for landraces can be considered.

In order to conduct association studies, the extent of LD and the decay of LD have a great influence on how to analyze association mapping and the SNP markers needed (Flint-Garcia et al. 2003). The results of LD showed that, the LD decay at a higher distance in genome D, than in genomes A and B. Genome B exhibits the lowest level of LD decay. Based on these results, fewer markers are required to detect target QTLs on genome D using GWAS than those required for detecting QTLs on the other genomes (Liu et al. 2017). A comparison of the SNP numbers for each genome reveals that, the D genome had the lowest number of SNPs followed by genomes A and B, respectively. Thus, it can be concluded that our SNPs and wheat population are suitable for GWAS analysis of traits related to target alleles. There is a high chance to identify target QTL with large and small effects based on the high and low LD found across the three genomes (Würschum et al. 2011). Other researchers have reported the same LD decay pattern across all three wheat genomes (Liu et al. 2017; Ayana et al. 2018). A large number of marker pairs were found in the B and A genomes whereas the younger D genome had a smaller number of markers. The same results were reported by others (Berkman et al. 2013; Edae et al. 2015). The higher diversity observed in the A and B genomes could be related to their older evolutionary background and due to gene flow from T. turgidum as opposed to lack of gene flow from Ae. tauschii to bread wheat (Dvorak et al. 2006; Jordan et al. 2015).

Totally 36 and 390 MTAs were significantly (P-value < 0.001) related to leaf rust resistance in Original and imputed datasets, respectively. However, only the relationship of the 19 high-confidence (FDR ≤ 0.05) SNPs across 12 chromosomes with previously identified *Lr* genes/QTL are explained below (Table 10) and the other SNPs are shown in tables 12 and 13. These markers represent 15 loci spread through chromosomes 1B, 2B, 3A, 3B, 4A, 5B, 5D, 6A, 6B, 6D, 7B, and 7D. The consensus map constructed by Maccaferri et al. (2015) was utilized to compare the significant SNPs identified in the study with previously cataloged *Lr* genes and QTLs. Figure 6 shows the schematic display of these resistance loci onto standardized chromosomes with similar length.

**Fig 6.**
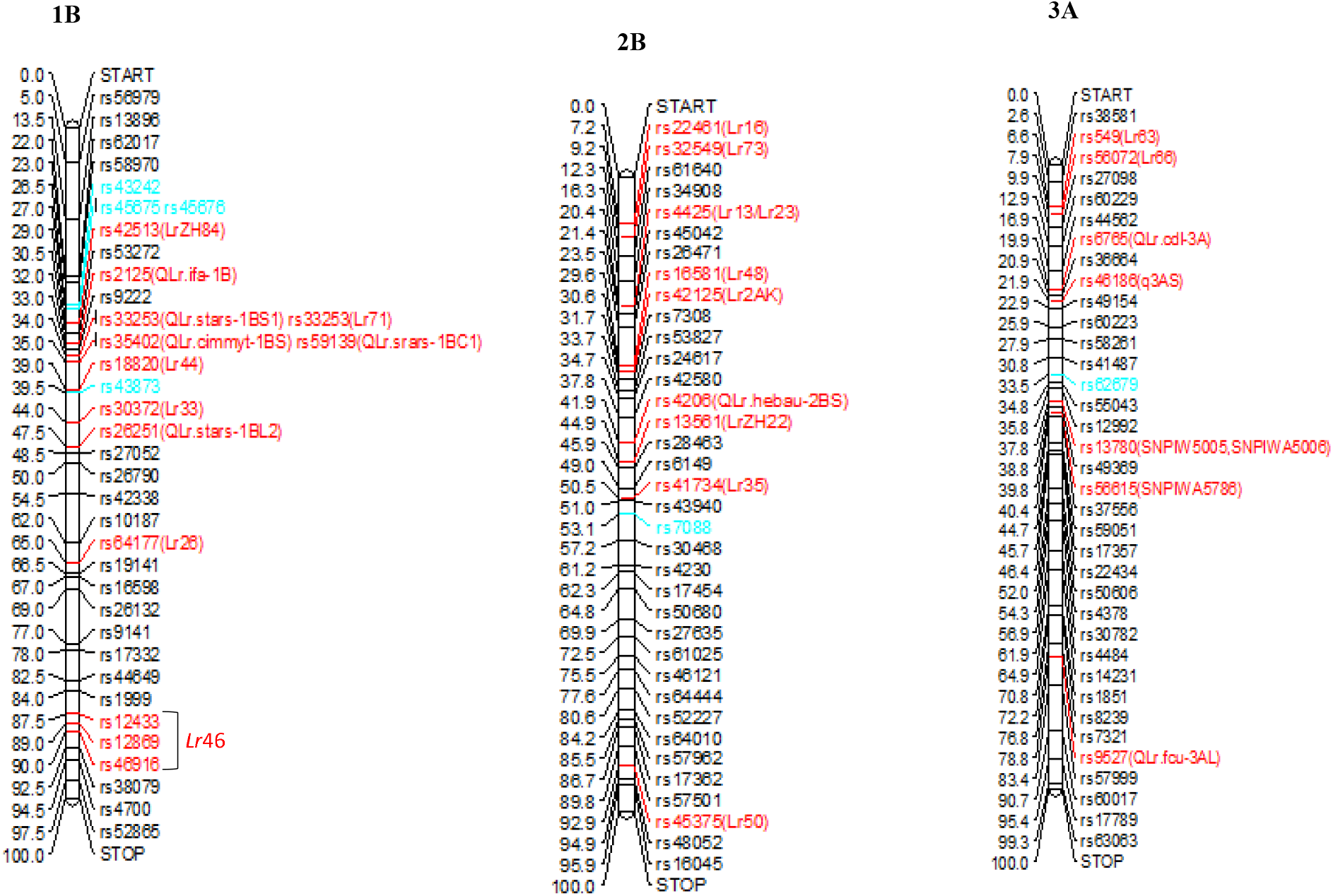
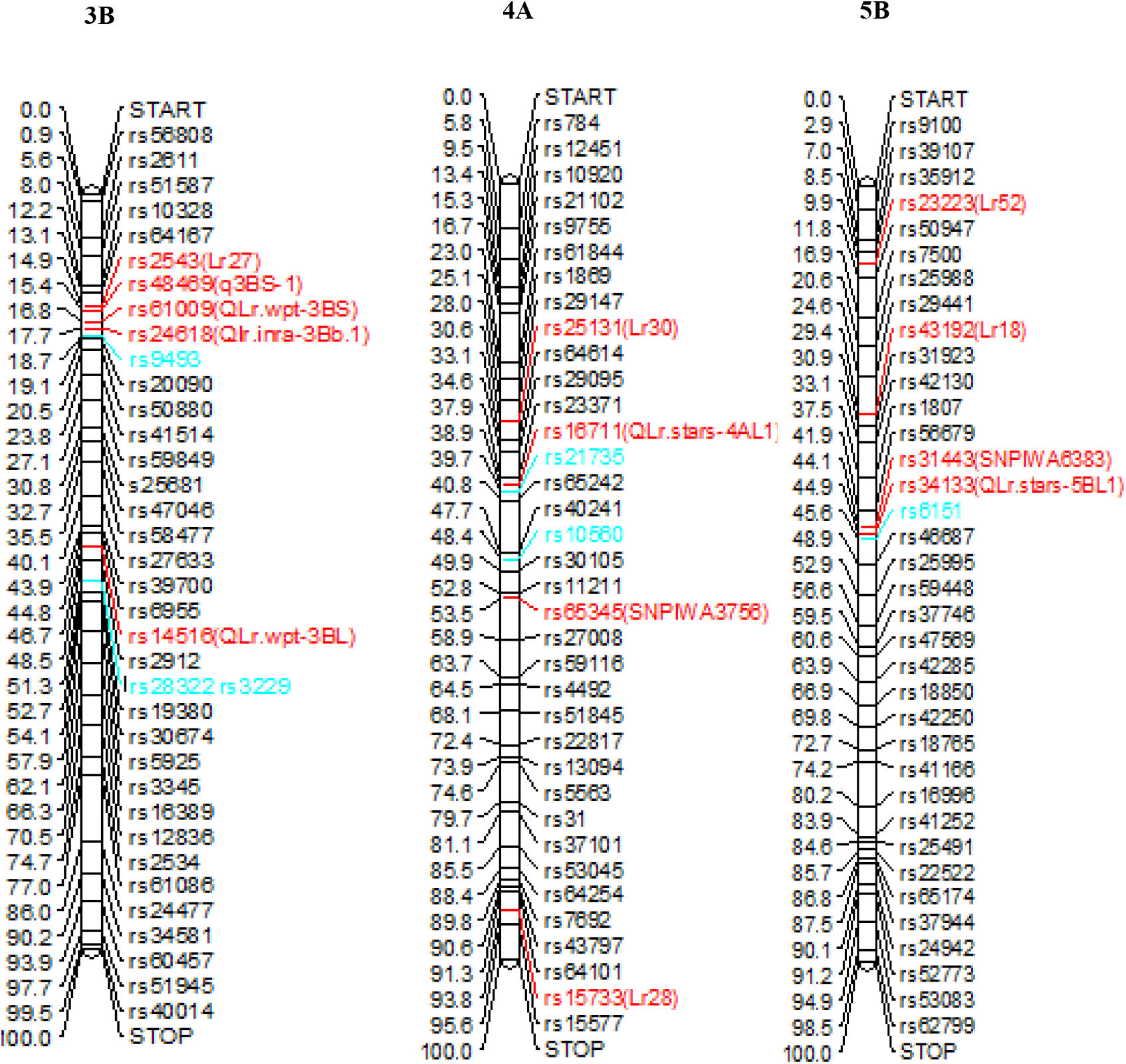
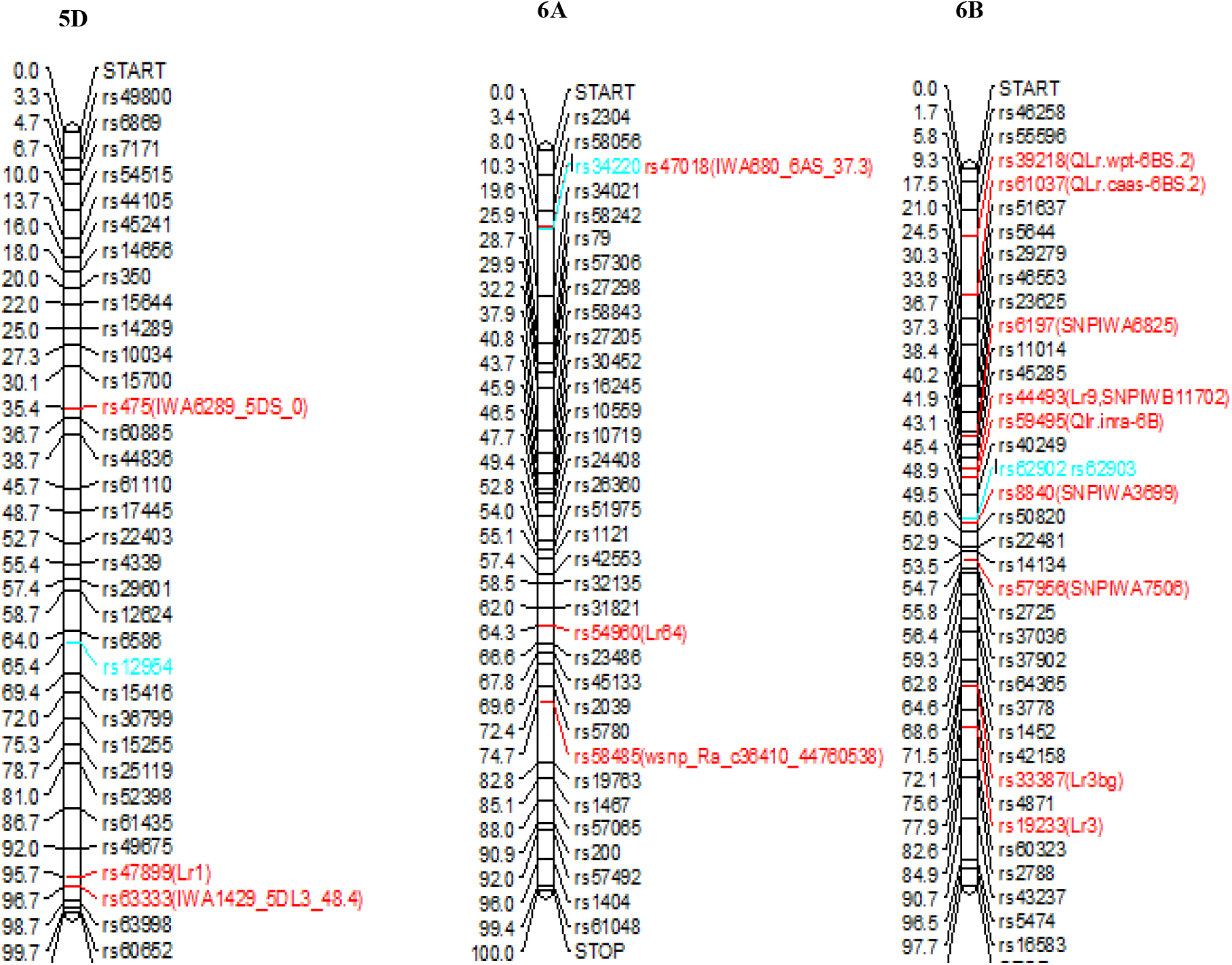
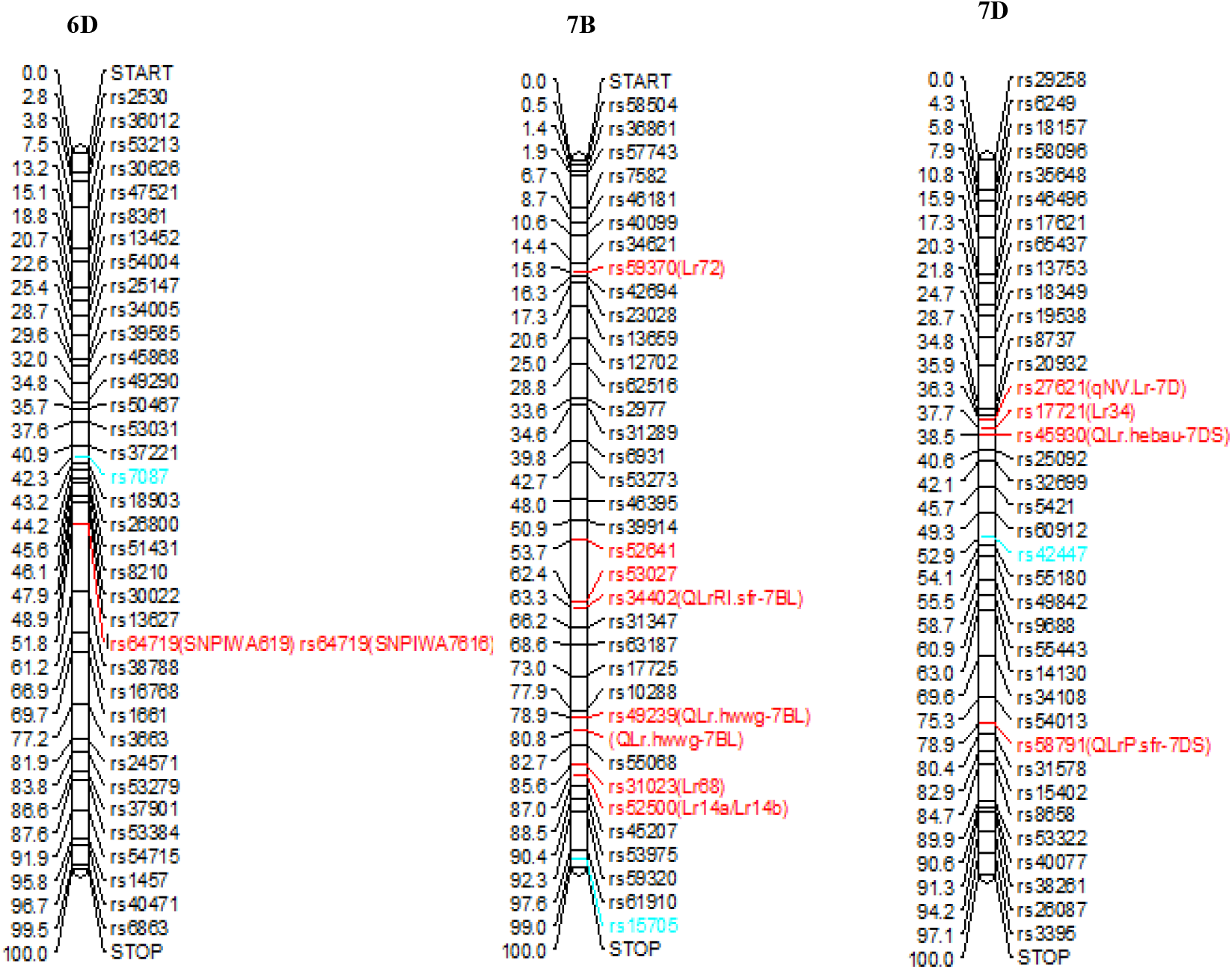
Chromosomal locations of quantitative trait loci (QTL) detected significant for resistance to leaf rust (LR) in this study relative to known *Lr* genes or QTL on those chromosomes based on the wheat consensus genetic map (Maccaferri et al. 2015). Markers detected significant for leaf rust resistance in this study are all in blue font and the previous detected genres/QTls are red fount. For better readability, not all markers are presented in this figure

### Chromosome 1B

GWAS identified four SNPs rs43242 (30.143cM), rs45675 (30.711cM), rs45676 (30.711cM) and rs43873 (45.006cM) for resistance to the PKTTS race. Nine known *Lr* genes, *Lr26, Lr33, Lr44* (Dyck and Sykes 1994), *Lr46* (Singh 1998), *Lr51* (Helguera 2005), *Lr55* (Brown-Guedira 2003), *Lr71* (Singh et al. 2013), *Lr75* (Singla et al. 2017), and *LrZH84* (Zhao et al. 2008), and five QTLs, *QLr*.*stars-1BC1* (Li et al. 2016), *QLr*.*cimmyt-1BS* (Rosewarne et al. 2012), *QLr*.*stars-1BS1* (Li et al. 2016), *QLr*.*ifa*-*1B* (Buerstmayr et al. 2014), *QLr*.*stars-1BL2* (Li et al. 2016), are mapped on chromosome 1B. Of these, *Lr26, Lr44, Lr51, Lr55*, and *Lr71* were originated from *Secale cereal*, spelta wheat, *Triticum speltoides, Elymus trachycaulis*, and spelta wheat, respectively (Dyck and Skyes 1994). Since no *Secale cereale, spelta*, and *Triticum speltoides* were involved on our GWAS panel, these markers are unlikely to be these genes. *Lr46*, from spring wheat cultivar CIMMYT Pavon 76 (Singh et al. 1998), *Lr75*, from wheat cultivar Arina (Schnurbsch et al. 2004), and *QLr*.*ifa-1B*, confer APR, and since the experiment was performed in the seedling stage, these markers are unlikely to be genes. *LrZh84*, probably derived from wheat cultivar Predgomaia, has been effective in the field for >30 years in China. Other QTLs mapped in this region, *QLr*.*stars-1BC1, QLr*.*cimmyt-1BS*, and *QLr*.*stars-1BS1* (Li et al. 2016), showed seedling resistance, and based on consensus map (maccaferi et al. 2015), three identified SNP markers (rs43242, rs45675, and rs45676) have almost same position with this QTLs, so it is likely these markers related to this QTLs. SNP marker rs43873 (45.006cM) were located close to QTL, *QLr*.*stars-1BL2* (Li et al. 2016). This QTL was mapped to be related to response leaf rust resistance in the seedling stage. Therefore, based on the genetic positions of the SNPs, it seems that they are probably associated with the previous QTLs.

### Chromosome 2B

Of the QTLs identified in GWAS in both datasets in FDR < 0.05, marker rs7088 on 2B at 59.184cM, was discovered to be related to resistance to the PFTTT race. *Lr* genes including *Lr13* (Dyck et al. 1966), *Lr16* (McCartney et al. 2005), *Lr23* (McIntosh and Dyck 1975), *Lr48* (Bansal et al. 2008), *Lr73* (Park et al. 2014), *LrZH22* (Wang et al. 2016), *LrA2K* (Sapkota et al. 2019), *Lr35* (Seyfarth et al. 1999), *Lr50* (Brown-Guedira 2003) and three QTL, *QLr*.*cimmyt 2BS* (Rosewarne et al., 2012), *QLr*.*hebau-2BS* (Zhang et al., 2017), and *QLr*.*uga-2BS* (Spakota et al. 2019), were also identified on chromosome 2B. Of these, *Lr13* (originated from Fontana), *Lr48* (Originated from CSP44), and *Lr35* are APR genes. As a result, the QTLs detected on chromosome 2B are unlikely to be APR genes. The other genes ie *Lr16, Lr23, Lr73*, and *QLr*.*cimmyt-2BS* are seedling resistance genes. Also, *Lr50* was derived from T. timopheevii armeniacum, since T. timopheevii armeniacum was included on our GWAS panel, these markers are unlikely to be this gene. According to the consensus map (Maccaferri et al. 2015), these genes are nearly co-located with these QTLs.

### Chromosome 3A

SNP rs62679 was identified at 57.65cM on chromosome 3A, which carries *Lr63* (Kolmer et al. 2010), *Lr66* (Marais et al. 2010) genes, and three QTLs, *SNPIWA5005, SNPIWA5006*, and *SNPIWA5786* (Kertho et al. 2015). *Lr63* and *Lr66* genes were derived from *Triticum monococcum* and *Aegilops speltoides*, respectively. Given that no *Triticum monococcum* and *Aegilops speltoides* were involved in our GWAS panel, these two genes are unlikely to be rs62679. SNP rs62679 was mapped near a previously mapped QTL, *SNPIWA5005, SNPIWA5006*, and *SNPIWA5786*, and therefore, the locus on 3A found in this study can be attributed to these QTLs.

### Chromosome 3B

On chromosome 3B, we identified SNPs rs28322, rs3229, and rs9493 that were related to seedling resistance to PKTTS, PKTTS, and PDKTT, respectively. Other *Lr* gene/QTLs that have been already reported close to rs28322, rs3229, and rs9493 include two *Lr* genes, *Lr27* (Mago et al. 2011) and *Lr74* (Bansal et al. 2014) and four QTLs, q3BS-1 (Li et al. 2016), *QLr*.*wpt-3BS* (Gerard et al. 2018), *Qlr*.*inra-3Bb*.*1* (Azzimonti et al. 2014), and *QLr*.*wpt-3BL* (Gerard et al. 2018), which in order to determine whether the SNP markers and previously identified genes/QTLs are related, further genetic analysis is required.

### Chromosome 4A

SNPs rs21735 and rs10560 at 62.15cM and 75.83cM, respectively, were detected in this research to be associated with resistance to PKTTS race in the seedling stage. Using the consensus map (Maccaferi et al. 2015) the SNP rs21735 is located in the same region as *QLr*.*stars-4AL1*. Therefore, it seems that rs21735 is probably associated with *QLr*.*stars-4AL1* (Li et al. 2016). Also, based on the consensus map, Marker rs10560 was identified in the vicinity of marker *IWA3756* (48.39cM). So, according to the consensus genetic map (Maccaferi et al. 2015) for both SNPs, it appears to be associated to previously identified QTLs, which are effective against the PKTTS race.

### Chromosome 5B

SNP rs6151, associated with PKTTS race in seedling stage, was observed near the genomic region of *QLr*.*stars-5BL1* (Li et al. 2016) and *IWA6383_5BL_138*.*8* (Turner et al. 2016). Furthermore, the *Lr18* (Carpenter et al. 2017) and *Lr52* (Hiebert et al. 2005) leaf rust-resistant genes, originated from *T*.*aestivum*, are mapped on the 5BL chromosome. Based on virulence/avirulence profile PKTTS, it has virulence on *Lr18* indicating that rs6151is unlikely to be *Lr18*. Based on the position of QTLs on the consensus map and their origin, the identified marker is likely related to QTLs, *QLr*.*stars-5BL1* and *IWA6383_5BL_138*.*8*.

### Chromosome 5D

SNP rs12954 was detected on chromosome 5D at 111.55cM. *Lr1* gene and two QTLs, *IWA6289_5DS_0* and *IWA1429_5DL3_48*.*4* are located on chromosome 5D (Turner et al. 2016; Gao et al. 2016). SNP rs12954 is effective for resistance against the PKTTS race. PKTTS race used in this study is virulent to the *Lr1* gene indicating that rs1294 is unlikely to be *Lr1* (Table 1). *IWA6289_5DS_0* is an APR QTL, confer slow rusting resistance, it is unlikely that SNP rs12954 is *IWA6289_5DS_0*. Also, SNP rs12954 was mapped far from *IWA1429_5DL3_48*.*4*, So that they are about 30 cM apart from each other. Therefore, it is likely that the genomic region tagged by SNP rs12954 related to a different QTL that confers resistance to leaf rust during seedling stage.

### Chromosome 6A

The SNP rs34220 was detected significant for resistance to leaf rust on chromosome 6A (Figure 5, 6; Table 10). Three catalogued *Lr* genes, *Lr56, Lr62* (Marais et al. 2008), and *Lr64* (Kolmer et al. 2010), and two QTLs, *IWA680_6AS* and *6A_t1* (Gao et al. 2016; Turner et al., 2016) have already been detected on chromosome 6A for leaf rust resistance. *Lr56, Lr62*, and *Lr64* are seedling resistance genes originated from *Aegilops sharonensis, Aegilops neglecta and, Triticum dicoccoides*, respectively (Somo et al. 2016; Kolmer et al. 2010). Due to the lack of genetic materials that carries these genes in our GWAS panel, it is unlikely that this locus represents *Lr56, Lr62*, and *Lr64. IWA680_6AS* and *6A_t1* (Turner et al. 2016; Gao et al. 2016), both QTLs identified on chromosome 6A confer APR. According to the genetic locus of gene/QTLs on the consensus genetic map and their origin, SNP identified in this position maybe associated with distinct loci for leaf rust resistance; however, more studies are needed to discover their associations between them.

### Chromosome 6B

On chromosome 6B, we identified two SNPs rs62902 and rs62903 in a same position (47.83cM), that were related to seedling resistance for PDKTT race. Other *Lr* genes and QTLs that have been previously identified close to these SNPs include Four known genes, *Lr3a, Lr3bg, Lr3ka* and, *Lr9* (McVey and Long, 1993) and, *6B_3, 6B_1, IWA7873, IWA7506, IWA5785, IWA8192, IWA6142, 6B_4, IWA3131, IWA3133, IWA5785, IWA6826, IWA6825, IWA7873, IWA8192, IWA6142, 6B_3, 6B_3, IWA596, IWA3699*, and *IWA7506* (Kertho et al. 2016) QTLs which requires further genetic studies to discover the association between the gene tagged by rs62903 and identified QTL/genes.

### Chromosome 6D

Marker rs7087 was mapped to the proximity of two previously mapped QTLs on chromosome 6D. According to the consensus map, the genetic map position of rs7087 (42.29) was 8.86 and 11.29 cM from *IWA619* and *IWA7616*, respectively (Kertho et al. 2016). Based on genetic map position, it is likely that rs7087 is correspond to *IWA619* or *IWA7616* leaf rust resistance QTLs. Further genetic research will be needed to found the association between rs7087 and previously identified QTLs.

### Chromosome 7B

Four previous identified *Lr* genes (*Lr14a, Lr14b, Lr68*, and *Lr72*) and a QTL, *QLr*.*hwwg-7BL*, (Lu et al. 2017) were already mapped on chromosome 7B within the region where rs15705 SNP was identified (Figure 6). Among the four *Lr* genes previously mapped on 7B, *Lr14a* and *Lr72* are from durum wheat (*T. turgidum diccocides*), and two genes, *Lr68* and *Lr14b* are from common wheat (McIntosh et al. 1995; Herrera-Foessel et al. 2012). *Lr68* is an APR gene and provides a high level of slow rusting resistance (Herrera-Foessel et al. 2012), this suggests that is it unlikely rs15705 corresponds to *Lr68*. Marker rs15705 is a SNP for resistance to PKTTS race and this race is virulent to the *Lr14a* gene indicating that rs15705 is unlikely to be *Lr14a*. Previously reported QTL for chromosome 7BL, *QLr*.*hwwg-7BL*, is an APR gene for leaf rust resistance (Li et al. 2014). Based on the relative length distance in consensus map (Maccaferri et al. 2015), the other QTLs detected on the 7BL >20cM are distanced from the detected marker. Further studies, such as utilize SSR markers for GWAS, allelism test or diagnostic marker analysis, can facilitate the determination of the association between rs15705 and reported gene/QTLs on chromosome 7BL.

### Chromosome 7D

Three known *Lr* genes *Lr19, Lr29*, and *Lr34*, and three QTLs, *qNV*.*Lr-7D* (Riaz et al. 2017), *QLr*.*hebau-7DS* (Zhang et al. 2017), and *QLrP*.*sfr-7DS* (Schnurbusch et al. 2004), were already mapped on chromosome 7D within the region where rs42447 was identified (Fig. 6). *Lr19, Lr29*, and *Lr34* were derived from *Thinopyron ponticum, Thinopyron ponticum*, and *T. aestivum* Terenzio, respectively. As no genetic material carrying *Thinopyron ponticum* was used in our GWAS analysis, rs42447 is unlikely to represent *Lr19* and *Lr29*. Likewise, three other QTL, QTL, *qNV*.*Lr-7D, QLr*.*hebau-7DS*, and *QLrP*.*sfr-7DS*, and *Lr34* gene are APR, therefore it is unlikely rs42447 be these gene/QTLs. As a result, rs42447 was found on chromosome 7D where no Pt resistance genes or QTLs had previously been identified. Therefore, the SNP rs42447 identified in genomic region 7D (83.31 cM) appears to be related to novel sources of resistance and could be valuable in breeding programs to enhance resistance to leaf rust.

Annotation of SNP sequences to the genes in *Triticum aestivum* L. proved our findings that these genomic regions encode proteins that are key components of signaling pathways that are activated in response to biotic and abiotic stresses. In general, these stresses change the expression of related genes in plants, for instance, increase or decrease of essential metabolites, changes in enzyme activity and protein synthesis, also the production of novel proteins (Zhu, 2016). For example, ATP binding protein (Lagudah 2011), ATPase activity (Heath, 1997), catalytic activity (Dmochowska-Boguta et al. 2015), carbohydrate-binding (Wu et al. 2020), nucleic acid binding (Zhang et al. 2019) were reported in earlier studies to be linked to plant diseases resistance. These genes are present in genomic regions associated with resistance traits and can be considered possible candidate genes for resistance against diseases as well as for future cloning of these loci.

## Conclusions

GWAS is an effective strategy for the discovery of molecular markers related to genes and QTLs in wheat. In this research, we assessed a diverse panel of 320 varieties and landraces of Iran for their response to five *Pt* races, PKTTS, PKTTT, PFTTT, PDTRR, and PDKTT, and have been detected ten wheat accessions highly resistant to all five *Pt* races. Totally, GWAS identified 19 QTL highly significant for resistance to leaf rust on chromosomes 1B, 2B, 3A, 3B, 4A, 5B, 5D, 6A, 6B, 6D, 7B, and 7D. Among these, a total of three SNP, on chromosomes 5D, 6A, and 7D, respectively, have been identified on genomic regions where no previously cataloged *Lr* genes has been reported from T. aestivum that represents potential novel loci for leaf rust resistance. Other significant SNPs, have been identified near known *Lr* genes or QTLs, and so, further research is required to approve the detected markers in this study to determine their relationship. These markers can be important targets for marker-assisted selection and fine mapping of functional genes after further validation.

